# A systemically delivered AAV-CFTR gene therapy for cystic fibrosis

**DOI:** 10.1101/2025.03.20.642115

**Authors:** Lindsey W. Plasschaert, Cian Stutz, Ester Otarola, Lionello Ruggeri, Rachel Valdez Misiolek, Arthur Nuccio, Jianyu Shang, Rayman Choo-Wing, Bruck Taddese, Arnaud Decock, Catherine Quigley, Katie Kubek-Luck, Isabelle Warnant, Daher Ibrahim Aibo, Maria Magnifico, Mirjam Buchs, Ge Tan, Louise Ashley, Hsu-kun Wang, Rebecca Watson, Amy Lin

**Affiliations:** Work was conducted at the Novartis Institutes for BioMedical Research, Cambridge, Massachusetts 02139, USA; Work was conducted at the Novartis Institutes for BioMedical Research, CH-4056 Basel, Switzerland; Work was conducted at the Novartis Institutes for BioMedical Research, San Diego, CA 92121, USA

## Abstract

Cystic fibrosis (CF) is the most common monogenic lung disease and results from mutations in the *Cystic Fibrosis Transmembrane Conductance Regulator (CFTR).* There have been over 2000 variants identified in patients that result in loss of function of the CFTR protein leading to systemic disease and respiratory failure in adolescence. While some variants encode proteins with residual activity that can be corrected or potentiated by CFTR modulators, at least 10% of CF individuals cannot tolerate the modulators or have nonsense mutations which fail to make any protein. For all people with CF, a mutation agnostic gene replacement strategy could provide a cure for CF lung disease. Here, we propose using a systemic route of administration to deliver a functional *CFTR* minigene cargo with a lung tropic AAV capsid. This would serve to reach multiple organs, most importantly the lung epithelium, and would provide a functional *CFTR* transgene that could be expressed in any cell type with a ubiquitous promoter. To achieve this, we generated the smallest *CFTR* minigene tested in an AAV delivery to date. We demonstrate its expression and function following transfection in cell-based assays and restoration of function in primary CF airway cells after viral delivery. Furthermore, we identify an AAV capsid that can transduce alveolar and airway epithelium with systemic delivery in non-human primates. These data provide tools for delivering a functional *CFTR* minigene that fits within the packaging capacity of an AAV and demonstrate lung transduction with an AAV following systemic delivery in a large animal model. This strategy first and foremost can reach target airway cells by circumventing the strong mucosal barrier in CF airways but may also provide a method by which to restore CFTR function in additional CF affected organs.

## Introduction

Cystic fibrosis (CF) is a life-threatening genetic disease that affects many tissues in the body, but the primary cause of morbidity is lung failure. It results from mutations in the *Cystic Fibrosis Transmembrane Conductance Regulator (CFTR)* which encodes an ion channel that localizes to the apical membrane of epithelial cells and is critical for anion transport across several mucosal tissues. Loss of CFTR-mediated ion transport in these tissues leads to dehydrated, sticky secretions. In the lung, this results in the adherence of mucus secretions harboring bacteria that can no longer be removed from the airway by mucociliary clearance. Chronic, recurrent infections remodel the lung and eventually reduce lung function. In addition to lung failure, pancreatic and gastrointestinal complications from CF lead to poor nutrient absorption and growth early in life. Aggressive, comprehensive care including enzyme therapy and antibiotics improved the lifespan of people with CF up to 30-40 years of age (McBennett et al., 2022).

Further extension of life in people with CF is attributed to the development of CFTR modulators, small molecules which bind to and improve the function of variant CFTR protein. CFTR potentiators, the first of which was made available in 2012, improve the gating of CFTR at the cell surface which regulates ion transport across the epithelial membrane. Another type of modulator, CFTR correctors, act on misfolded protein and aid in its trafficking to the apical surface. CFTR potentiators and correctors have been combined into a triple combination therapy (elexacaftor/tezacaftor/ivacaftor) that demonstrated improved lung function in people with *ΔF508*, the most common *CFTR* variant (Heijerman et al., 2019; Middleton et al., 2019). The available CFTR modulators can theoretically improve lung function in more than 90% of people with CF who have variants that make dysfunctional protein. However, people with Class 1 *CFTR* variants which do not make any protein due to unstable or prematurely degraded mRNA (approximately 6-10%) and people unable to tolerate modulators for various reasons (possibly as high as 20%) are underserved by the existing CFTR modulators (Hubert et al., 2017). This population would greatly benefit from a *CFTR* gene therapy. In addition, a gene transfer strategy which delivers a functional copy of the *CFTR* gene could be adopted by people with any *CFTR* variant.

Following the cloning of the *CFTR* gene in 1989, which identified the genetic cause of CF, it immediately became a promising application for gene therapy (Riordan et al., 1989). Since respiratory failure is the leading cause of death in people with CF, a lung directed gene therapy would provide the most benefit in preventing morbidity. Over the next three and half decades, *CFTR* gene therapies have been tested with a variety of delivery vectors and administration routes and formulations in clinical trials (reviewed in Plasschaert et al., 2024). *CFTR* gene therapies have been delivered via bronchoscope or nebulization directly to the lung. Viral and non-viral delivery vectors including adenovirus, adeno associated virus (AAV), and lipid nanoparticles (LNP) have shown promise in transferring *CFTR* DNA or mRNA to the airway epithelium, but have failed to demonstrate sufficient gene transfer to result in an improvement in efficacy as measured by percent of predicted forced expiratory volume in one second (ppFEV1) (Alton et al., 2016; Harvey et al., 1999; Rowe et al., 2023; Wagner et al., 2002). While adenovirus and LNPs have proven to be more challenging in their current iterations due to increased immunogenicity, AAV has demonstrated enormous potential as a gene delivery vector.

The initial *CFTR* gene therapy clinical trials utilized AAV2 and its internal promoter to express *CFTR* (Wagner et al., 2002). Poor tropism for apical surface receptors of the airway epithelium and weak promoter activity of AAV2 are largely blamed for insufficient CFTR gene transfer and expression. Additionally, a limiting factor for AAV as a gene delivery vector is its inability to package genomes over 4.7 kilobases (kb). To this end, AAV-based *CFTR* gene therapies have attempted to shorten both the *CFTR* gene and strong, ubiquitous promoter sequences in order to generate a cargo that can be packaged effectively (Ostedgaard et al., 2002, 2005). Finally, while a lung-directed administration implies best access to the lung, there is evidence that the mucus film lining the airways could block transduction of target epithelial cells, and that mucus plugging could limit the distribution of an inhaled aerosol (Carneiro et al., 2020; Duncan et al., 2018). Furthermore, systemically (intravenously) administered gene therapies have demonstrated safety and efficacy in diseases of peripheral tissues (Al-Zaidy et al., 2019; Potter et al., 2021). Because CF affects many organs in addition to the lung, a systemically delivered gene therapy that can reach the lung, but also access other CF affected tissues would be the best option for people that cannot use modulators. To this end, we have generated a *CFTR* minigene small enough to fit within AAV genome and shown that it restores chloride transport in primary human airway epithelial cells. Additionally, we have demonstrated that an AAV capsid is capable of transducing airway and lung epithelium following systemic delivery in non-human primates. Together, these tools address an unmet need and provide a possible solution to widespread *CFTR* restoration in people with the most severe variants of *CFTR*.

## Results

### Structure-guided engineering of shortened CFTR

Due to the large size of *CFTR*, packaging of the full-length wildtype gene (4440bp) together with even a short promoter and polyA signal will push the packaging size limitations of AAV (Dong et al., 1996). Many efforts have been made by numerous labs to attempt to reduce the size of *CFTR* while maintaining its wildtype function (Carroll et al., 1995; Fischer et al., 2007; L. Zhang et al., 1998). Truncated forms of CFTR suffer from strongly reduced functional activity with one of the exceptions being a construct where 52 amino acids have been deleted at positions 708-759 in the regulatory domain (R-domain) (Ostedgaard et al., 2002).This construct, henceforth described as ΔR, has been shown to maintain wildtype function in various functional assays (Excoffon et al., 2024; Ostedgaard et al., 2005b, 2011; Steines, Dickey, Bergen, Excoffon, et al., 2016) and is currently being tested in the clinic (NCT05248230). Yet, despite the reduction of CFTR by 52 amino acids, the cargo size remains at the limit of what can be efficiently packaged and delivered by AAV which may lead to genome fragmentation and reduced viral titers. Therefore, we set out to engineer a novel truncated variant of CFTR with further reduced size.

Most of the published minigenes were engineered over 20 years ago when protein structural information was not yet available. Here, we aimed to capitalize on the availability of the more recently published high-resolution cryoEM structures of CFTR (Liu et al., 2017) and take a structure-guided approach combined with existing knowledge about previously tested deletions to engineering novel minigenes. Analysis of the CFTR protein structure led to six groups of deletions we wanted to explore (Figure 1). In the first group containing the extracellular region, we shortened the protruding extracellular loops (ECL). Special care was to avoid introducing any structural strain on transmembrane helices connected by these loops. The second group contained truncations of the N-terminal region. Several N-terminal deletions have been reported in the literature which included the deletion of one or several transmembrane (TM) domains (Δ-S118M, Δ259; Δ1-264 (Carroll et al., 1995; Fischer et al., 2007)). We could not observe any functional activity for those when transfected into HEK 293 cells (data not shown) and thus focused on N-terminal deletions preceding the first TM domain. The third group contained constructs where intracellular loops (ICL) were shortened. As with the ECLs, we ensured that shortened loops would not lead to structural constraints on the domains they were connecting. This included analysis of the open and closed forms of the CFTR structure (Z. Zhang et al., 2018). In the fourth group, we looked at shortening loops within the nucleotide binding domains (NBDs). NBDs are strongly conserved across orthologues and species and not many opportunities could be identified for truncation besides one extended loop region. The fifth group contained C-terminal deletions. Here we tested a previously described deletion (Zhang et al., 1998). Additionally, we designed a novel deletion which retained the acidic cluster previously described to impact CFTR function (EETEEEVQ, Ostedgaard et al., 2003) (Figure 1A). In the sixth group, we explored novel deletions in the R domain. In 2002, Ostedgaard et al., reported the impact of deletions of various parts of the R domain on the Cl^-^ current in airway epithelia. Except for the previously mentioned ΔR (deletion of amino acids 708-759), all deletions had major impact on bumetanide-sensitive short-circuit currents and/or basal currents. We hypothesized that for some of the deletions, the reason for the malfunction may not stem from the deletion of the particular sequence of amino acids but rather from fusing the two resulting ends and thereby introducing structural strain on the CFTR molecule. Thus, using molecular modeling, we calculated the minimal number of amino acids that the R-domain should contain to bridge the structural gap between positions M645 and N825. Based thereon we replaced the R-domain deletions with glycine-serine linkers of appropriate lengths with the aim to enable deletion of certain R-domain stretches without introducing structural strain (Figure 1B).

**Figure 1:**
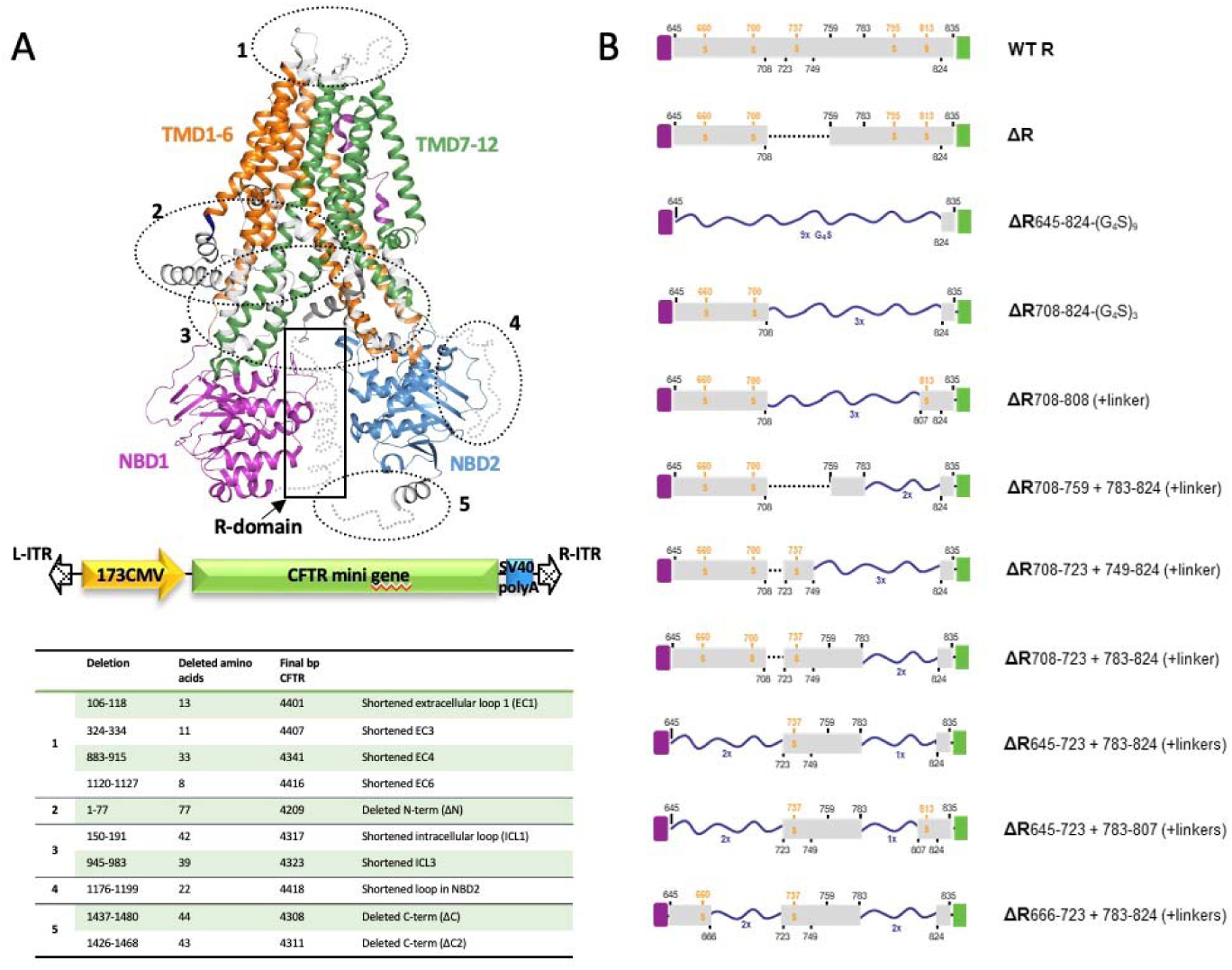
Structure-guided engineering of shortened CFTR constructs. (A,B) Cryo-EM structural analysis and knowledge of the functional protein domains modified in previous CFTR minigenes iterations led to shortening of 1) the extracellular loops (EC), 2) N-terminal domain (ΔN), 3) intracellular loops (ICL), 4) nucleotide binding domain (NBD), 5) C-terminal domain (ΔC) and 6) Regulatory domain (ΔR;**B**). These constructs were then cloned downstream of 173CMV promoter ((Ostedgaard et al., 2005b) in between the AAV2 ITR sequences and including an SV40pA tail (**A**). Illustration of the sixth group, in which R domain deletions were bridged by a series of glycine-serine linkers in order to prevent structural strain (**B**).

The >40 novel minigenes with structure guided engineering of unique deletions and linker sequences were transfected into HEK293T cells which do not express endogenous *CFTR*. Minigene expression was driven by a shortened CMV promoter previously described (173CMV; (Ostedgaard et al., 2005; Figure 1A) and screened for CFTR function using a FLIPR membrane potential assay. In response to forskolin stimulation, ion transport resulted in a change in membrane potential observable by increased luminescence. While many constructs with deletions in the N-terminus, ECLs and ICLs showed no signals, several minigenes carrying deletions in the C terminus and regulatory domains as well as one minigene that included a deletion in the nuclear binding domain demonstrated increased luminescence similar to full length (FL) CFTR (Figure 2A). We next assessed more sensitive and quantitative parameters of CFTR activity on a single cell basis using the patch clamp technique. Once again, HEK 293 cells were transfected with the *CFTR* minigenes or *FL CFTR*. While all minigenes tested displayed an ability to respond to forskolin stimulation, the amplitude of the current measured at +100mV was significantly reduced in minigenes containing an N terminal and intracellular loop 3 (ICL3) deletion (Figure 2B). We proceeded to test viral delivery and function of the minigene containing deletions in the C terminal and R domains (ΔR+ΔC) because it was the shortest transgene that retained similar function to *FL CFTR* in response to forskolin activation.

**Figure 2:**
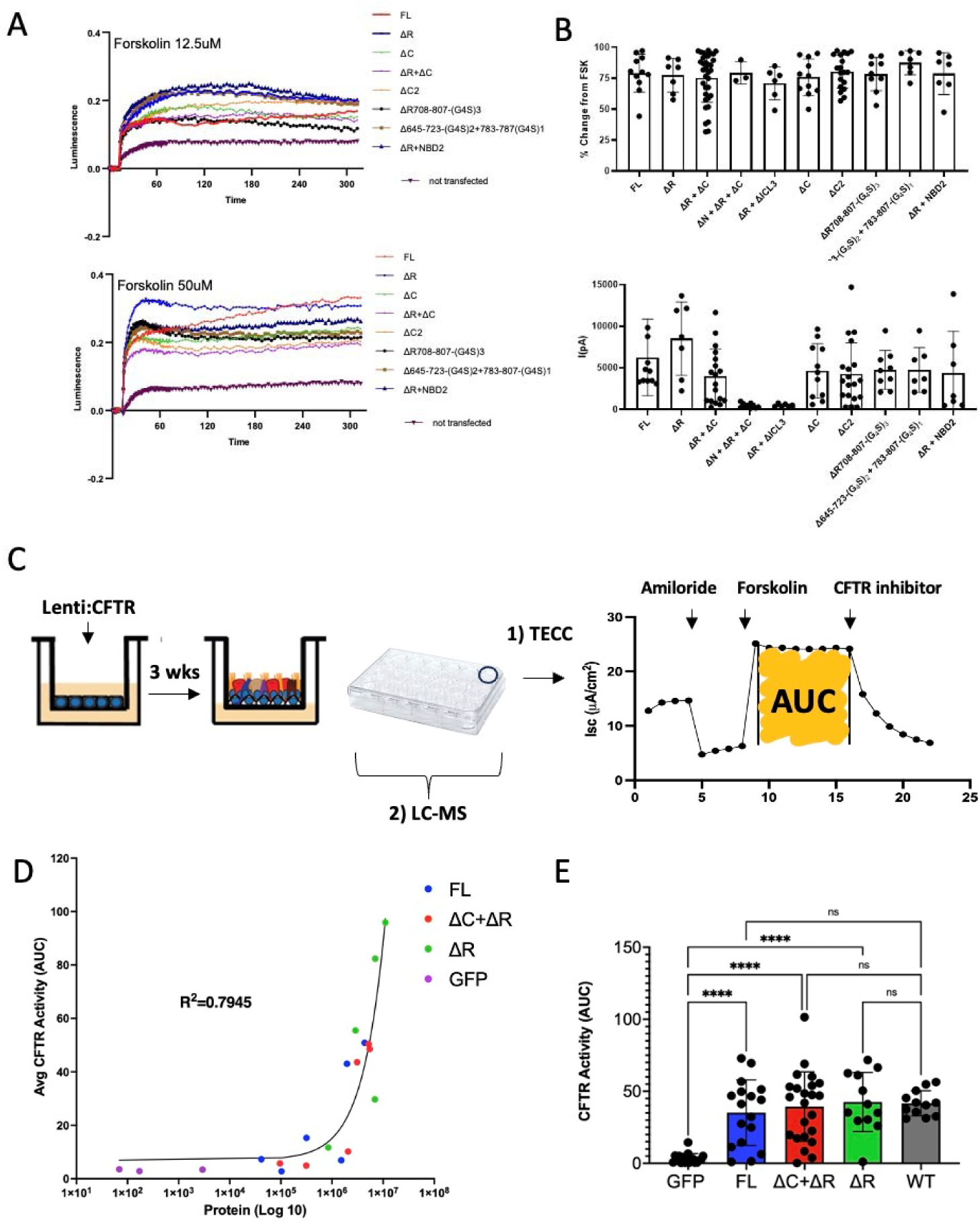
Functional analysis of CFTR constructs in cell-based assays demonstrates ion transport mediated by CFTR ΔR+ΔC is comparable to wildtype CFTR. (A,B) CFTR constructs were transiently transfected into HEK293T cells and stimulated with forskolin. CFTR activity was analyzed by luminescence due to changes in membrane potential using FLIPR assay (**A**) and for ion flow by automated patch clamp technique (**B**) (**C**) Next, primary human CF bronchial epithelial cells were seeded on transwell inserts and transduced with lentivirus expressing CFTR constructs. Following differentiation, ion transport (Isc; short circuit current) in individual transwells (n=6 per condition per experiment) was measured using the transepithelial chloride conductance assay (TECC) and the area under the curve (AUC) between forskolin addition and CFTR inhibition was quantified. Subsequently, cultures treated with identical lentivirus were pooled for protein extraction (resulting in n=1 peptide measurement per condition per experiment) and liquid chromatography mass spectrometry (LC-MS) was used to measure CFTR peptide and graphed against average CFTR activity for that condition (**D**). CFTR activity for each minigene following comparable protein delivery by lentiviral vector was measured and compared to GFP vector or untreated wildtype HBECs (**E**). (Two donors each for WT and CF HBECs were tested in 2 independent experiments; ****p<.0001)

The gold standard for measuring CFTR function in human airway cells preclinically is the Ussing chamber or transepithelial chloride conductance assay (TECC) performed on primary human bronchial epithelial cells (HBECs) grown at an air-liquid-interface (ALI). This electrophysiology assay has been used extensively to measure changes in CFTR channel function after potentiation or correction of mutant CFTR protein using modulators. Furthermore, restoration of Cl^-^ current in this preclinical assay translated to lung function improvement (ppFEV1) in clinical trials (Keating et al., 2018). In order to compare the activity of the ΔR+ΔC CFTR to FL CFTR in HBECs, constructs were packaged in lentiviral vectors capable of packaging larger transgenes than AAV, and therefore, could be used to test wildtype FL CFTR. We also included the ΔR minigene as a positive control as it is a well-studied minigene that has demonstrated similar activity to wildtype protein in preclinical assays. Lentivirus expressing GFP was included as a negative control and to confirm HBEC transduction by lentivirus. Lentivirus was delivered to CF HBECS containing Class 1 mutations which generate severely truncated protein or lack protein translation due to nonsense mediated decay of mRNA. Flow cytometry for GFP and electrophysiology for CFTR activity were performed three weeks later, after cultures had differentiated and polarized. CFTR activity was quantified as the area under the curve (AUC) of the current (Isc), between the time of forskolin stimulation and CFTR-172 inhibition (Figure 2C). Since functional titer of lentivirus can vary due to size of cargo and intracellular processing, we also wanted to determine how much CFTR protein was delivered with each lentivirus. Therefore, we optimized a light chromatography mass spectrometry (LC-MS) proteomics assay, a sensitive analytical method which allowed us to quantify CFTR peptide regardless of truncation status by measuring a specific peptide found in all three CFTR constructs. Individual wells of CF HBECs transduced with lentivirus expressing *GFP* or *CFTR* constructs were first measured for CFTR activity by TECC. Next, up to six wells were pooled for protein collection and quantification by LC-MS (Figure 2C).

We first plotted the relationship between average CFTR activity measured in TECC assay and amount of protein delivered by lentivirus and found a significant positive correlation between increasing protein and activity (Figure 2D; R^2^=.7945). Since there appeared to be a higher functional titer of lentivirus achieved with packaging of ΔR and thus a variable distribution of protein delivered with each lentivirus, we binned the samples by protein and analyzed the conditions in which more than one construct was represented and delivered protein was greater than endogenous protein measured in wildtype cultures (Figure S1A; bins 1-3). This analysis revealed similar activity between ΔR+ΔC and full length CFTR confirming it can transport chloride as efficiently as wildtype CFTR. Class 1 CF HBECs transduced with GFP did not induce CFTR activity, as expected. All three CFTR constructs had significantly increased CFTR activity compared to GFP and were not significantly different from CFTR activity generated in wildtype HBECs, thus demonstrating that the ΔR+ΔC CFTR minigene can stimulate vectoral ion transport in primary human airway epithelial cells (Figure 2E).

Having demonstrated that the engineered ΔR+ΔC CFTR minigene could restore CFTR ion transport in primary human CF airway cells when delivered with lentivirus, we wanted to determine whether it was sufficiently small to be packaged effectively within an AAV. Therefore, the ΔR+ΔC CFTR minigene was packaged in an AAV relevant for lung transduction and the product was run on a denaturing gel and sequenced to determine the percentage of DNA fragments at the correct size (Figure 3). Both the gel and sequencing data indicate that the majority of the DNA species are the appropriate size representing the ΔR+ΔC CFTR construct (4789 nt) demonstrating that the ΔR+ΔC CFTR minigene is sufficiently small to be packaged in an AAV.

**Figure 3:**
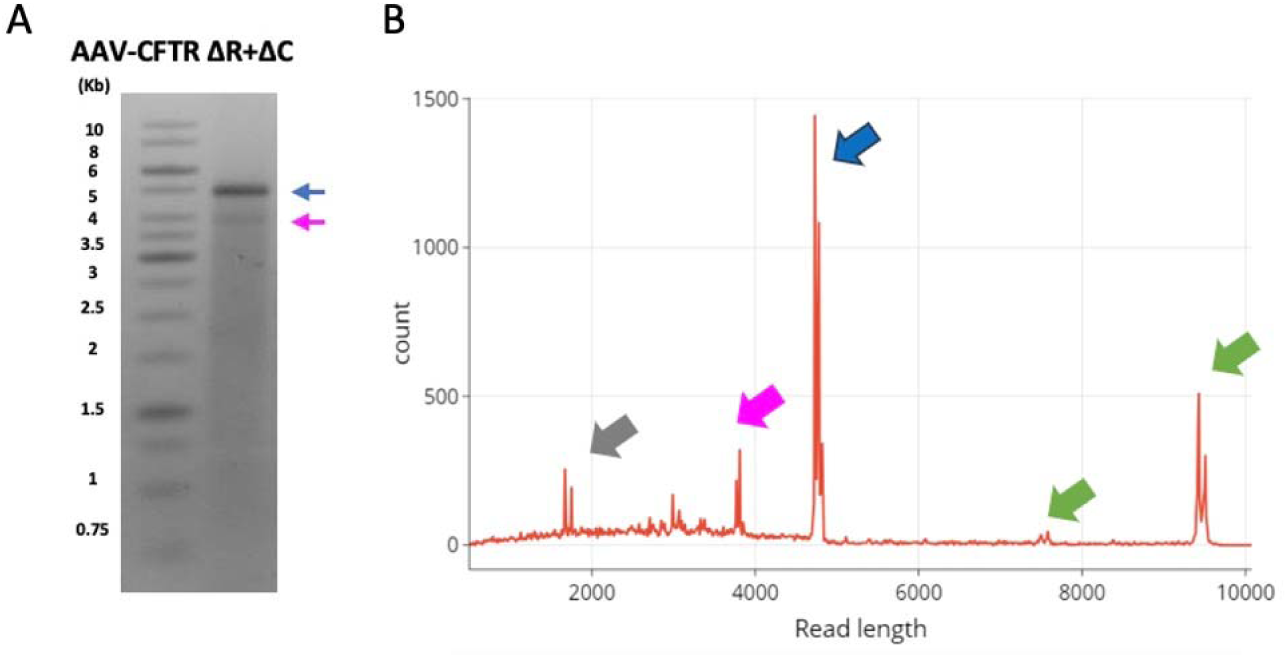
CFTR ΔR+ΔC can be packaged efficiently in an AAV with short 173CMV promoter. (a) ΔR+ΔC CFTR was packaged in an AAV with a shortened *(173)CMV* promoter. Vector genome integrity analysis by denaturing alkaline agarose gel (**A**) and vector genome integrity analysis by PacBio sequencing (**B**) was performed. Blue arrows represent full size of vector genome (4789nt). Pink arrows represent reverse packaged full plasmid backbone (3632nt) Green arrows represent the synthesized complementary strand called natural complement formation during amplification. Left arrows indicates the double sized backbone and right arrow indicates the double sized full vector genome. Grey arrow indicates partial vector genome containing ITR, 173CMV promoter and particle ΔR+ΔC CFTR sequence.

### AAVs transduce human airway basal cells by basolateral administration

We next wanted to determine if an AAV capsid could transduce human bronchial epithelial cells at sufficient levels to rescue CFTR-mediated ion flow across the epithelial sheet, a function critical for mucus hydration and mucociliary clearance. To this end, we first determined what percentage of cells need to express CFTR to restore wildtype activity. Previously, it has been reported that CFTR correction in 5-10% of cells in an epithelial sheet can restore normal activity (Johnson et al., 1992). Furthermore, it’s been reported that viral transduction of 20-25% of CF mutant cells with full length *CFTR* can restore normal CFTR activity (Excoffon et al., 2009; L. Zhang et al., 2009). We found in cell mixing experiments that cultures with 10% non-CF HBECs restore 5% of CFTR activity, cultures with 25% non-CF HBECs restore 30% CFTR activity and cultures with 50% non-CF HBECs are not significantly different from 100% non-CF cells (Figure S1B,C). The CFTR modulator combination lumacaftor/ivacaftor restored 14% of CFTR activity in vitro and resulted in a 2-4% improvement in ppFEV1 in clinical trials (Van Goor et al., 2011; Wainwright et al., 2015). Therefore, our data support previous findings that rescuing between 10-25% of the CF epithelium can restore CFTR activity to a level that will result in a clinically meaningful improvement in CF lung function.

A significant obstacle to delivering a CFTR gene therapy via inhalation is the concentrated, sticky mucus that accumulates in the absence of CFTR function. Alternatively, a gene therapy vector could be delivered intravenously to access the lung via the circulatory system, thus circumventing the mucosal barrier. However, this requires transduction by basolateral receptors, which has not been thoroughly assessed. We next sought to demonstrate that an AAV capsid is capable of transducing 10-25% of human airway epithelial cells by our intended administration to the basolateral rather than apical surface. To this end, we tested a variety of AAVs with known tissue tropism (Halbert et al., 2001). We delivered AAVs at 1 × 10^5^ vg/cell to HBEC cultures grown at ALI, via the apical side, mimicking an inhaled delivery, or via the basolateral side, mimicking a systemic delivery. We then assessed the transduction of basal cells vs. apical cells by flow cytometry using the basal cell markers Integrin Alpha-6 (ITGA6) or Nerve Growth Factor Receptor (NGFR) (Figure 4). From a panel of AAV serotypes, we confirmed two were efficient at transducing HBECs and performed a dose response with these AAVs expressing an mcherry cargo with a full length CMV promoter at MOIs of 5 × 10^4^, 1 × 10^5^ and 5 × 10^5^ vg/cell. These capsids were more efficient at transducing basal cells by basolateral delivery than they were at transducing apical cells by apical delivery suggesting they have preference for basolateral receptors (Figure 4B). One capsid, “AAVLungVar,” achieved ∼40% transduction of basal cells with basolateral delivery, while the other achieved approximately half of that (20%) consistent with prior comparisons between lung tropic AAVs (Limberis et al. 2009). These data indicate that an AAV capsid can transduce the total population of HBECs at levels sufficient for restoration of CFTR-mediated ion transport.

**Figure 4:**
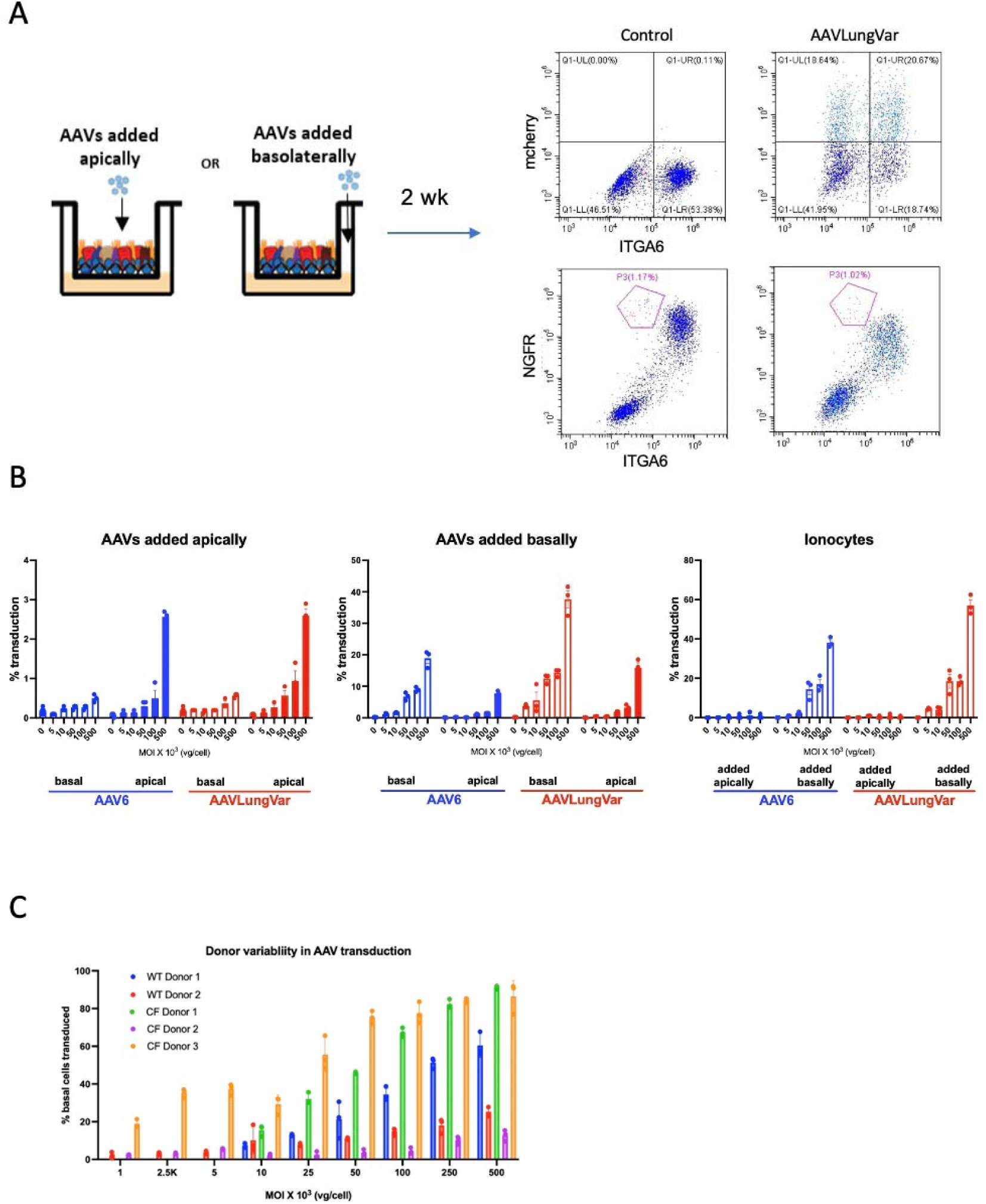
AAVs transduce human bronchial epithelial cells and ionocytes via basolateral delivery in vitro. **(A)** AAVs expressing mCherry cargo driven by CAG promoter were added either apically to primary human bronchial epithelial cells (HBECs) grown in transwells to mimic lung-directed delivery or basolaterally in well media to mimic a systemic delivery route. Following differentiation, cells were trypsinized and stained for markers ITGA6 and NGFR to identify apical (ITGA6-/NGFR-), basal (ITGA6+/NGFR+) or ionocyte (ITGA6-/NGFR+; P3) cell populations. Sample shown is from basolateral delivery of AAVLungVar at 2.5 × 10^5^ vg/cell **B**) Increasing concentrations of AAVs demonstrated a dose response in transduction of each cell population. **C**) AAVs were added basolaterally to different donor derived HBECs (n=2 non-CF; 323353 and 429581 and n=3 CF; KKD017K, KKD043N, KKD023N) to compare basal cell transduction by mCherry flow cytometry.

We also asked whether the pulmonary ionocyte, a cell type we discovered to be a high expressor of CFTR, was transduced (Montoro et al., 2018; Plasschaert et al., 2018). Ionocytes are highly NGFR positive and ITGA6 negative, so we were able to identify them in the sorting panel we used for basal cells (Figure 4; Population 3). In contrast to the total apical population (including secretory cells and ciliated cells), ionocytes were more highly transduced by basolateral delivery (Figure 4B). In fact, the AAVLungVar transduced 60% of the rare ionocyte population at the highest MOI delivered basolaterally. Given the morphology and basolateral projections of ionocytes previously described, we speculate that ionocytes may have better access to or preferential receptors for transduction on the basolateral domain. We further assessed transduction of basal cells from WT and CF cultures using basolateral delivery and found differences in transduction from donor to donor that was not dependent on CFTR status (Figure 4C).

### AAVLungVar transduces airway epithelial cells *in vivo* in mouse and non-human primate

We sought to demonstrate that a lung tropic capsid could transduce airway epithelium following systemic delivery, with the potential advantage of also transducing additional CF affected organs by this route of administration. Thus far, we had demonstrated certain AAV serotypes could transduce human airway epithelial cells via the basolateral domain and there is evidence that AAV serotypes can transduce mouse airway by intratracheal delivery *in vivo* (Limberis et al., 2009; H. Wu et al., 2024). Additionally, it’s been reported that AAV tropism and biodistribution can vary depending on route of administration (Konkimalla et al., 2023; Zincarelli et al., 2008). Therefore, we compared mouse lung transduction by three AAV serotypes, including the AAVLungVar that potently transduced HBECs by basolateral delivery *in vitro*. Each contained a tdtomato cargo and was delivered by either inhalation (oropharyngeal aspiration) or intravenously (tail vein injection). The AAVLungVar was superior at transducing proximal conducting airway and distal airway epithelium at a dose of 5 × 10^12^ vg/kg by inhalation. In fact, we saw little to no transduction of distal airway by an alternative AAV at this dose (Figure 5A). Though we detected robust transduction by AAVLungVar with inhaled delivery, there was no overlap between the basal cell marker cytokeratin 5 (KRT5) and tdtomato suggesting inhaled delivery did not target mouse basal cells (Figure 5C). We systemically delivered a three-fold higher dose (1.5 × 10^13^ vg/mouse) of the AAVLungVar and an AAV shown to have broad biodistribution by tail vein delivery in mouse (Figure 5B). These data confirmed that a broadly tropic AAV (“AAVBroad”) in mouse was superior at transducing lung by tail vein delivery than the AAVLungVar that was found to be potent in primary human airway cells. These data support previous findings that route of administration can alter transduction efficiency for AAV serotypes (Furusho et al., 2024; Konkimalla et al., 2023).

**Figure 5:**
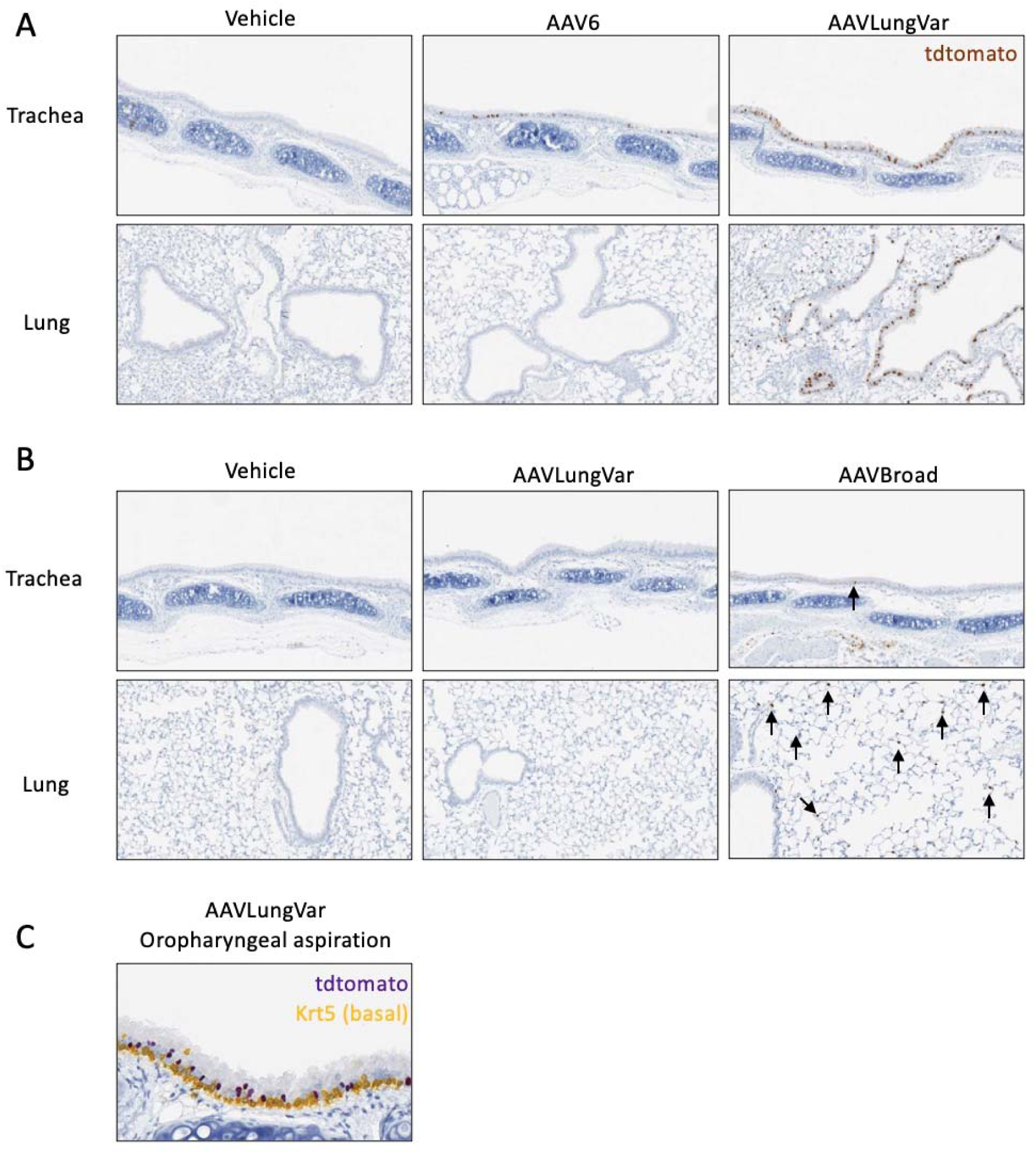
AAVs demonstrate different lung tropism depending on route of delivery in mouse. AAVs expressing tdtomato reporter driven by ubiquitous CAG promoter were delivered to mice (n=8 per AAV) by oropharyngeal aspiration (OA) at a dose of 5 × 10^12^ vg/kg (**A**) or by tail vein delivery at a dose of 1.5 × 10^13^ vg/kg (**B;** arrows mark epithelial cells transduced by systemic AAV delivery). Tracheas (proximal airway) and lungs (distal airway) were collected two weeks following AAV dosing and counterstained for tdtomato using an RFP antibody. Tracheas from AAVLungVar OA-treated animals were further analyzed by costaining for tdtomato and the basal cell marker cytokeratin 5 (KRT5; **C**).

Thus far, we had demonstrated that the AAVLungVar capsid could transduce human airway epithelial cells by basolateral delvery *in vitro* but demonstrated low tropism for mouse airway by systemic delivery *in vivo*. Species-specific differences in AAV transduction and cargo expression have been demonstrated and can result from changes in expression of AAV receptors, altered chromatin accessibility, and other factors that are still being elucidated (Batista et al., 2020; Das et al., 2022a; Gonzalez-Sandoval et al., 2023). Non-human primates represent the most similar anatomy and immune response to humans, both of which may impact transduction of an AAV, particularly with systemic delivery. Therefore, we dosed cynomolgus macaques with 1 × 10^14^ vg/kg intravenously with the AAVLungVar and additional capsids tested in mouse expressing an mCherry reporter under control of the 173CMV promoter, the same promoter used to drive the virally delivered CFTR minigenes. We measured viral genome DNA in every tissue and performed IHC for mCherry protein in select tissues (n=2 monkeys per group). As expected, virus biodistribution to the liver was higher than any other tissues for all capsids, and no significant differences could be derived between the three capsids given the low animal number (Figure S2A). Interestingly, the liver IHC demonstrated that AAVLungVar transduced liver endothelial cells, while the other two capsids transduced hepatocytes by morphology (Figure S2B). Strikingly, only AAVLungVar transduction led to lung expression of mCherry in ∼5-10% of the bronchial surface epithelium and ∼80% of the distal airway alveoli (Fig 6). There was no mCherry protein detected in lung epithelial cells from primates treated with the other two vectors. Co-hybridization of *mcherry* mRNA with cell type-specific markers by RNAscope revealed that <1% of the basal airway epithelial cells were *mcherry* positive (Figure S3). These data demonstrate that the AAVLungVar is capable of transducing the airway epithelium by systemic delivery. Transduction of surface epithelium and submucosal glands was 5-10%, an amount suggested to be sufficient to make meaningful improvements in CFTR-mediated ion homeostasis and CF lung disease (Johnson et al., 1992; Ramalho et al., 2002); however, transduction of basal stem cells is low (<1%). Since turnover of these cells would result in dilution of AAV over time, redosing would likely be required to maintain sufficient CFTR activity long term.

**Figure 6:**
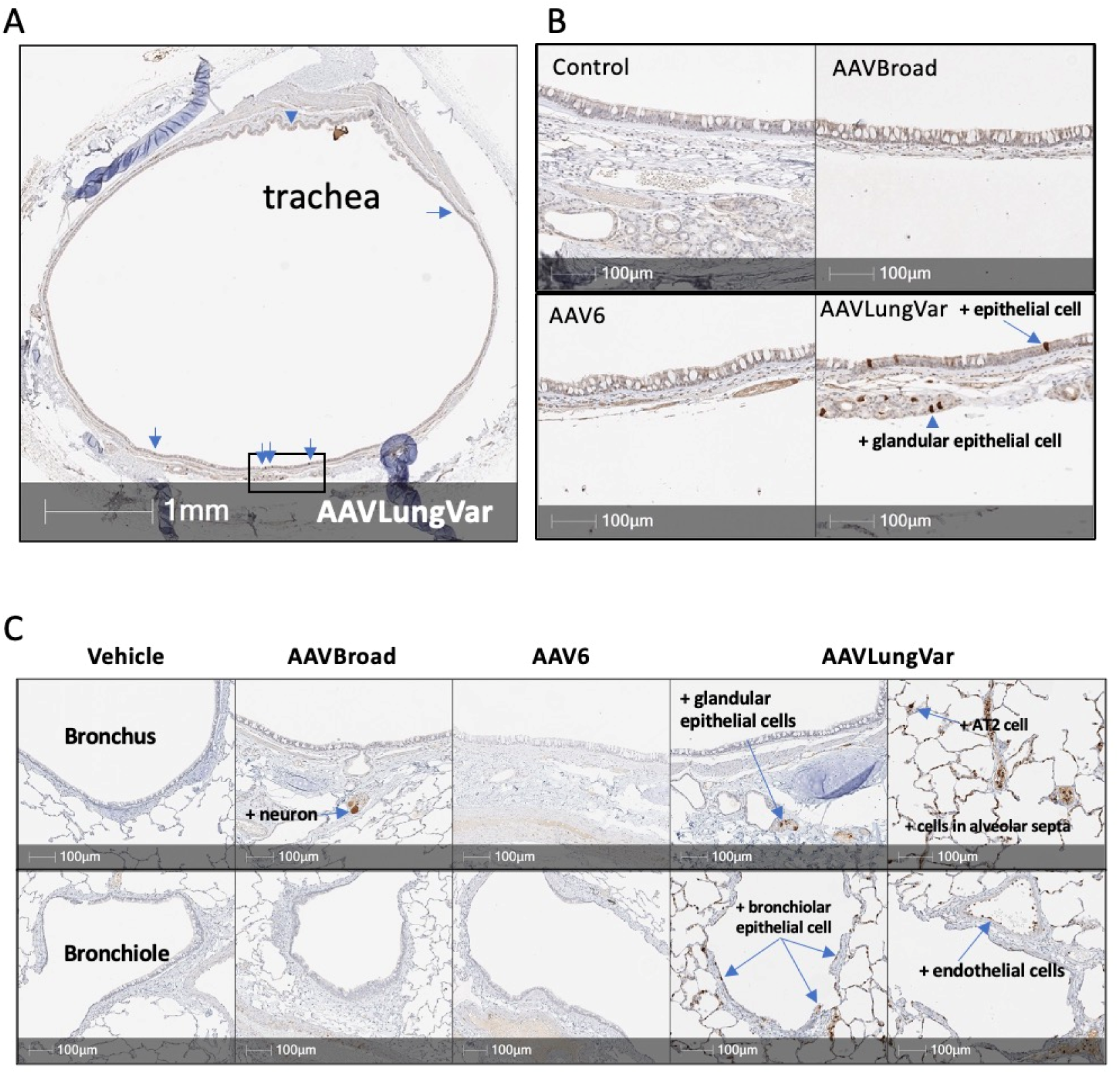
Immunohistochemistry images indicating that AAVLungVar transduces airway epithelium of non-human primate (NHP) following systemic intravenous (IV) delivery. AAVs expressing mcherry cargo driven by a ubiquitous, shortened 173CMV promoter were delivered to NHP (n=2 per AAV) at a dose of ∼1 × 10^14^ vg/kg by intravenous injection. Tissues were harvested three weeks post dosing and stained for mCherry using an mCherry antibody. Representative images of trachea (**A,B**) and lung from proximal to terminal bronchioles (**C**) demonstrate that AAVLungVar transduced upper airway epithelium, submucosal gland cells, alveolar epithelial cells and terminal bronchiolar epithelial cells and endothelial cells.

## Discussion

Since the discovery of the CF disease causing mutation in 1989, there have been more than 40 clinical trials testing *CFTR* gene therapies (clinicaltrials.gov). Despite this, there is still not an approved *CFTR* gene therapy. Several reasons for lack of clinical efficacy have been explored (Cooney et al., 2018; Plasschaert et al., 2024). Poor gene transfer by delivery vectors may be due to low affinity or tropism for airway cells or an inability to access appropriate receptors through concentrated mucus. Immunogenicity against certain vectors such as adenovirus and LNPs has been demonstrated to limit durability of gene expression. Finally, AAV vectors, which are thought to be relatively less immunogenic, are restricted in their ability to package larger genomes such as the full length *CFTR* coding region. Here we developed the tools for a *CFTR* minigene that could be packaged in an AAV and systemically delivered to bypass the mucosal barrier and transfer *CFTR* to multiple tissues, including and most importantly, the lung epithelium.

Engineering cargos that are small enough to fit within AAVs has been a challenge for multiple disease indications and successful strategies include shortening the target gene or splitting cargos between multiple AAVs (Hirsch et al., 2016; Wang et al., 2000). In preclinical development, functional demonstration of the engineered cargo in primary human cells is necessary when not expressing the wildtype transgene. We furthered previous efforts to shorten the *CFTR* gene for an AAV vector by combining the ΔR and ΔC deletions. The resulting CFTR construct functioned similarly to full length CFTR in both cell-based assays that assessed ion flow through the channel, as well as in primary human airway cells, the gold standard for CFTR therapies, which assessed ion flow across an epithelium (Figure 2E). It is important to note that CFTR has many functional domains involved in its trafficking, gating, regulation, and binding interactions with other proteins. Therefore, novel CFTR minigenes are best tested *in vivo* in order to demonstrate rescue within its physiological context where it functions in surface hydration, mucociliary clearance and airway surface liquid pH regulation (Plasschaert et al., 2024). Testing CFTR gene therapies in the context of Class 1 mutations, as was done here, is also important as CFTR has been shown to dimerize, and a well-studied minigene Δ1−264 CFTR was able to rescue CFTR variants via transcomplementation but was not sufficient to transport chloride on its own (Cebotaru et al., 2013). The ΔR+ΔC minigene is a smaller derivation of CFTRΔR which has been tested *in vitro* and *in vivo* is currently being tested in the clinic (Ostedgaard et al. 2002).

We were able to demonstrate function of ΔR+ΔC and packaging efficiency with a shortened CMV promoter in an AAV capsid. Next to be determined is the functional output of the ΔR+ΔC minigene generated here following delivery with a capsid such as the AAVLungVar. However, considerations for functional rescue by lentivirus versus AAV must be taken into account. For example, lentivirus is integrated into the genome while AAV is not, which could impact the level of expression. Additionally, the tropism for certain cell types, differentiation and polarization of those cells and their ability to employ exogenous CFTR could impact expression and function. While it has been shown that proteasome modulators such as doxorubicin can improve expression of AAV cargos in airway epithelial cells (Yan et al., 2004; L. N. Zhang et al., 2004), we were able to detect transduction by AAV-mCherry by flow cytometry without the use of doxorubicin.

CFTR antibodies have significant background and nonspecific staining for flow cytometry and immunohistochemistry (IHC) (Sato et al., 2021). Therefore, in this study, we utilized LC-MS method to measure CFTR protein, which was able to detect small amounts of CFTR and would be best suited for measuring CFTR delivery by an AAV. Furthermore, the method allows differentiation between single amino acid changes relevant for comparing various minigenes or measuring species-specific CFTR delivery to a non-CF animal model. Therefore, this method could be a valuable tool for assessing gene transfer of *CFTR* gene therapies preclinically and clinically. Expression of the cargo within an AAV capsid can also be impacted by epigenetic factors adding to the complexity of AAV delivery and cargo expression in different cellular contexts (Das et al., 2022; Gonzalez-Sandoval et al., 2023). Therefore, preclinical studies should measure expression of the lead therapeutic candidate in a relevant animal model as it may differ from expression of a reporter cargo.

Previous clinical trials utilized a direct lung administration for gene therapies for CF. While the lung offers a direct route of administration, the mucosal and epithelial immune barrier has evolved to protect against entry of pathogens such as viruses and lipid-coated membranes (McCarron et al., 2023). In the last five years, several AAV-based gene therapies have been approved with an intravenous route of administration, two of which, Zolgensma and Elevidys, are intended to treat non-liver peripherally located tissues. These trials and approvals demonstrate the overall safety and efficacy of delivering an AAV systemically at doses that can reach peripheral tissues without overt liver toxicity. To our knowledge, this is the first study to examine transduction of lung epithelial cells in NHP by systemic AAV delivery. In fact, we demonstrate that AAVLungVar, a vector shown to be lung tropic in mouse and human airway epithelial cells, is capable of transducing NHP trachea and alveolar epithelium, while other capsids did not. There did not appear to be large differences in viral genome distribution between the three capsids in any tissue, emphasizing the need for assays measuring expression of the cargo in tissues and cell types of interest. Interestingly, AAVLungVar appeared to transduce endothelial cells preferentially to hepatocytes. It is yet unclear whether this would alter the high liver absorption or liver toxicity of high doses of AAVs (Hordeaux et al., 2024), although those may also be mitigated by using a cell type specific promoter (rather than CMV) to restrict expression.

Species specific differences in AAV tropism has made preclinical testing of AAV gene therapies challenging and may necessitate testing in multiple species (Watakabe et al., 2015). Furthermore, evolved capsid screening in a single animal model can result in selection of species-specific transduction that does not translate to humans (Steines, et al., 2016). Here, we tested capsids via multiple routes of administration in primary human airway cells, mouse, and NHP by systemic delivery to demonstrate cross-species translatability. Interestingly, AAVLungVar showed high transduction of primary human airway cells and NHP airway cells by systemic delivery but was less potent than AAVBroad in mouse. Results such as these may necessitate using surrogate capsids in species-specific disease models and testing clinical capsids in the most human relevant models for safety and toxicology. Additionally, we found differences in transduction profiles depending on the route of administration (Figure 5) as demonstrated previously (Konkimalla et al., 2023). This necessitates testing the clinical capsid via a relevant route of delivery in the best preclinical animal model in order to understand safety associated with biodistribution and expression.

While we did identify a capsid that could transduce NHP airway with systemic delivery, AAVLungVar did not transduce the proximal airway or basal stem cells at levels that would be sufficient to maintain durable CFTR expression in the airway. A method to improve durability may include a more direct bronchial artery injection to improve transduction by allowing the gene therapy to reach the lung prior to the liver. Alternatively, a genome editing or gene insertion approach may be sufficient for long term expression even if it reaches a small percentage of progenitor cells. The tools evaluated here move us toward a systemically delivered CF gene therapy with the potential to transduce multiple organs. Novel delivery vectors with improved targeting to lung such as SORT-LNPs or targeted AAVs are being developed (Cheng et al., 2020; Goertsen et al., 2022). However, specific vector targeting to lung or restricted expression via a lung-specific promoter may leave people with CF comorbidities in other organs. Ultimately, a gene therapy aimed at restoring CFTR function systemically, such as the approach used here, would greatly benefit all people with CF, and those with severe Class 1 variants in particular.

## Methods

### Structure-based CFTR minigene engineering

CryoEM structures (PDB ID: 5UAK and 6MSM for dephosphorylated and phosphorylated CFTR, respectively) served as the foundation for structural analysis and molecular modeling. These analyses were conducted using MOE v2020.09 (Molecular Operating Environment, Chemical Computing Group ULC, Montreal, QC) and BioLuminate v2021 (Schrödinger, LLC, New York, NY). Additionally, Alphafold 2.2.2 (PMID: 34265844) was employed for the structural prediction of CFTR minigenes. Potential deletions with minimal impact on CFTR structure and conformational changes were identified through visual inspection. Here special attention was given to existing knowledge around domains and motifs and to maintaining the integrity of secondary structure elements. Deletions were made such that fusion of the two resulting ends would not result in the morphing of remaining secondary structure elements. Furthermore, movements of secondary structure elements and domains between the dephosphorylated and phosphorylated structures were considered, ensuring that deletions would not interfere with structural rearrangements. Models were generated by constructing fusions between deleted residues and assessing the impact on geometry. The optimal glycine-serine (gly-ser) linker length for bridging R-domain deletions was determined based on the structure of dephosphorylated CFTR. The structure indicated that the R domain inhibits NBD dimerization, with residues 825–843 of the R domain inserting into an opening between TMD9 and TMD10. To calculate the minimum number of amino acid residues necessary to connect the structured regions (M645 to N825) without requiring adjustment, thereby maintaining low strain energy. Strain energy for various lengths of alanine and 4GS linkers was calculated using BioLuminate’s Cross-Linking Proteins protocol. This process identified both the minimum residue length—the approximate number of alanines required to form a suitable linkage—and the maximum residue length corresponding to the wild-type sequence length. Based on these calculations, gly-ser linker lengths were chosen accordingly, depending on the number of residues deleted from the R-domain.

### Construction of CFTR minigene cargo plasmids

DNA fragments with internal deletions for the newly designed CFTR variants were gene synthesized at Geneart AG (Regensburg, Germany). Some variants of the CFTR gene were generated by PCR using Gibson assembly (Gibson DG, Young L, Chuang RY, Venter JC, Hutchison CA, Smith HO. 2009) including N- and C-terminal deletions with appropriate DNA primers obtained from Microsynth AG, Balgach, Switzerland. Final fragments were ligated into an cis-plasmid vector in which a truncated version of the cytomegalovirus promoter (173CMV (Ostedgaard et al., 2005b)) drove cDNA expression. An SV40pA (Choi et al., 2014) element followed the CFTR gene. Correct insertion and final sequences were confirmed by Sanger sequencing (Microsynth AG). Final plasmids were transformed in TOP10 Chemically Competent *E. coli* (Ref. C404010, Invitrogen) and DNA preps were prepared using “Qiagen Plasmid Plus Maxi” kit (Ref#12963). Integrity of L- and R-ITRs was tested by restriction digestion with XmaI (Ref. R0180L, NEB). Endotoxin levels were measured using Endosafe with a cutoff of <0.5EU/mg.

### CFTR membrane potential assay

CFTR minigenes were transiently transfected using standard conditions into HEK293T cells in suspension in serum-free V3 medium two days before experiment. Transfected HEK293T cells were treated with membrane potential dye for 30 minutes at 37 °C then loaded onto fluorometric imaging plate reader (FLIPR) where forskolin ranging from 100-.0006uM was injected to activate CFTR. Subsequent fluorescent changes were recorded for 6 minutes and the formula dF/F = [Ft- (mean of Fb/Fb)] was plotted using Graphpad. Fb = average of the first 10 measurements before injection of forskolin. Membrane potential kit “blue” was used (Molecular devices, #R8034).

### Cell preparation for Qpatch

Cells were grown in T150 tissue culture flask and minigenes were transfected 48hrs before study. On the day of the experiment, cells grown to ∼70% confluency were rinsed once in 10 mL of sterile DPBS and the cells were lifted with 3mL of Detachin (Amsbio, AMS.T100100) for ∼5 min. When cells were dislodged from culture flask surface, 3mL of CHO-S-SFM (Thermo scientific, 31033020) with 25mM of HEPES (Gibco, 15630) was added and aspirated. Aspirated cells were centrifuged at 300g for 5 min and resuspended at 3 million cells/mL in CHO-S-SFM with 25mM HEPES.

### Qpatch

Whole-cell patch-clamp experiments) were performed on a QPatch-48 automated electrophysiology platform (Sophion Biosciences) using the 48-channel, single-hole QPlate (resistance 2.0 ± 0.4 MΩ). The external buffer (in mM) was 160 NaCl, 4.5 KCl, 2 CaCl2, 1 MgCl2, 10 HEPES, 5 glucose, pH 7.3 The internal buffer (in mM) was 130 KCl, 2.7 CaCl2, 0.88 MgCl2, 10 EGTA, 10 HEPES, 2 Na2ATP, pH 7.2. 20μM Forskolin (Sigma, F3917-10mg) was used as a CFTR activator and 30 μM CFTRinh-172 (Sigma, C2992) as an inhibitor. The voltage protocol applied for CFTR recordings was determined by a holding potential at-40mV to test a ramp between-100 to +100mV in 1 second. This voltage protocol was applied every 10 s during the liquid protocol is applied. Recordings were performed at room temperature (20–25°C). The liquid protocol applied was: 1) External solution (20 voltage protocol run), 2) external solution with DMSO 0.3% (25 voltage protocol run), 3) Forskolin (60 voltage protocol run) and 4) CFTRinh-172 (60 voltage protocol run). The QPatch software automatically generates sweep traces and current time (IT) plots. General quality control (QC) filters: 200MΩ seals, <100 pA leak current, current window for max outward current 50pA<x<5nA, Whole cell capa citance <4pF, Rs >50MΩ. The IT plots were generated from the currents at +100mV.

### Flow cytometry

Cells were rinsed with warm DPBS 1X (Gibco, 14190144), then collected using 0.05% Trypsin EDTA (Thermo Fisher, 25300054) and pelleted at 300g for 5 min. Cells were then suspended in 2% FBS DMEM with EDTA and filtered through a 40-μm strainer before being analyzed by flow cytometry or cell sorting. Cells analyzed by flow cytometer were spun down and resuspended with staining buffer BSA (BD Bioscience, 554657). Antibodies NGFR-APC 1:200 (BioLegend, 354108) and ITGA6-PE 1:500 (BioLegend, 313612) were used.

### Lentivirus production

Lentiviral packaging 4×10˄6 HEK293T cells were seeded in a 100-mm poly-d-lysine-coated dish (Corning BioCoat, 356469) one day before transfection with 14 ml of cell growth medium (DMEM (Thermo Fisher, Cat# 11965092), 10% FBS (Clontech 631106), 2mM l-glutamine (Invitrogen 25030), 0.1mM MEM Non Essential Amino Acids (Invitrogen 11140), and 1mM sodium pyruvate MEM (Invitrogen 11360)). For transfection, 7 μg of packaging plasmid DNA (ViraPower lentiviral Packaging Mix, Thermo Fisher K497500) was mixed with 5 μg of expression construct DNA and 36 μl Fugene6 (Promega, E2691). OptiMEM (Thermo Fisher, 31985062) was then added the mixture to a total volume of 800 μl. 293T cells were incubated with the transfection reagent mixture for 24 h before the growth medium was refreshed. At 72 h after transfection, the virus was collected, and filtered with 0.45µM PVDF Steriflip filter HAWP04700. Lenti-X™ GoStix™ Plus (631280) was used to detect viral titer. If the titer is low, Lenti-X™ Concentrator (631232) was used to concentrate the virus. Lentivirus was aliquoted and frozen at-20°C for future experiments. Packaged virus was added to HBEC cultures 1 h after cell seeding and then removed at feeding the following day.

### Human bronchial epithelial cell culture

Primary human bronchial epithelial cells (HBECs) from wildtype donors were obtained from Lonza (CC-2540; Lot, 323353, 429581) and CF donors were obtained from UNC tissue core (KKD023N, KKD029O; KKD017K; KKD043N). P0s were expanded twice with growth medium (500 ml BEGM medium (Lonza, CC-3171), 1 SingleQuots kit (Lonza, CC-4175)) in T75 flasks. After expansion, HBECs were seeded on 24-well Transwell plates (Corning, 3460) at a density of 44,0000 cells per Transwell. The cells were cultured in differentiation medium (250 ml BEGM medium, 250 ml DMEM medium (Thermo Fisher, 11965092), 1 SingleQuots kit) on both apical and basal sides of Transwells for the first 7 days. Then, medium was removed from apical side, and cells were cultured for another two weeks at an air liquid interface (ALI) condition. Cells were used for analysis after culture at ALI for 14 days and no more than 28 days. In HBEC studies analyzing tropism of AAV capsid serotypes, AAVs were added after 2 weeks of differentiation at ALI. In HBEC studies measuring CFTR activity from lentiviral vectors, vectors were added 1h after seeding to allow for transduction by lentivirus.

### Short-circuit current (Isc) measurements in Transepithelial chloride conductance (TECC) assay

For TECC studies, HBECs were cultured in 24-well transwell plates (Corning) at a seeding density of 44,000 cells per well. Tranwell plates containing differentiated HBECs were then bathed in Buffer (Kreb’s Ringer solution; 400 ml H2O, 25 ml 2.4M NaCl, 25 ml 0.5M NaHCO3, 25 ml 66.6 mM KH2PO4 + 16.6 mM K2HPO4, 25 ml 24 mM CaCl2 + 24 mM MgCl2, 0.9 g dextrose). Amiloride (Sigma, A9561) was added apically at 10 μM to inhibit Na+ absorption, then forskolin (Sigma, F6886) was added apically at 20 μM to stimulate cAMP and finally, CFTR-172 (Sigma-Aldrich, C2992) inhibitor was added apically and basally at 30 μM. Under these conditions, cAMP-stimulated Isc due to addition of forskolin could be attributed to CFTR-mediated Cl− secretion from basolateral to apical solution.

### Liquid Chromatography Mass Spectrometry

Following differentiation at air-liquid interface (ALI) in 24-well plates, HBECs were pooled from at least 6 wells for a given condition and dissociated using 0.05% trypsin–EDTA (Thermo Fisher, 25300054). Cells were then pelleted at 300g for 5 min, resuspended in PBS, then pelleted again. Supernatant was removed and cells were frozen at-80.

### Proteomics Sample Preparation

Sample pellets were processed using PreOmics iST kits, but with a few modifications. PreOmics iST lysis buffer with 1% Bond-Breaker TCEP was added to each pellet and heated at 95°C for 10 minutes and shaken at 1000rpm. A VialTweeter UP2006t sonicator was then used to lyse the cells and shear DNA. Protein concentration was determined with Pierce BCA assay kits. 50ug of lysate was used for PreOmics Trypsin/Lys-C digest. Before digest, 2.5fmol of heavy target peptide (Biosynth; Gardner, MA, USA) was spiked into solution. Enzymatic digest was performed at 37°C for 3 hours. After digest, the reaction was quenched for 15 minutes by adding TFA to a final 1% v/v concentration. The reaction vial was then centrifuged for 15 minutes at 10,000g to pellet the precipitated detergent in the PreOmics lysis buffer. The supernatant was saved for high pH fractionation. The High pH fractionation was performed using a Pierce High pH reversed phase fractionation kit. The manufacturer’s instructions were followed, but only Fraction 1 was retained since it contained both CFTR peptide targets.

### Liquid Chromatography Mass Spectrometry Targeted Analysis

Parallel reaction monitoring (PRM) targeted analysis was performed using a Thermo Orbitrap Eclipse (Thermo Scientific) with an EASYnLC 1200 (Thermo Scientific) in technical triplicates for WT Untransduced samples and technical duplicates for all other samples. 250ng lysate with 13amols of heavy peptide were directly injected onto a 25cm x 75µm Aurora Ultimate column (C18, 1.7µm; IonOpticks). The mobile phase was (A) 0.1% Formic Acid (LC-MS Grade; Honeywell) and (B) 80% Acentonitrile, 0.1% Formic Acid (Optima LC-MS grade; Fisher Scientific). Gradient elution at 400nL/min and 50°C was performed as follows: 0-5%B for 5 minutes, 5-13%B for 13 minutes, 13-100%B for 10 minutes, and 100%B for 8 minutes. The heavy and light peptide targets VADEVGLR and ACQLEEDISK, common to all minigenes and full length CFTR, were HCD fragmented with 19 and 21 NCE, respectively.

### PRM Data Analysis

Peptide intensities were extracted using Skyline v22.2.0.527 (University of Washington; Seattle, WA, USA). Intensities were normalized using the heavy spike-in. The quantification results were based off the more intense VADEVGLR peptide due to missing transition values from the less intense peptide ACQLEEDISK.

### AAV production

AAVs were produced by 3-plasmid co-transfection in HEK293 cells containing an adenoviral helper plasmid, a Rep-Cap plasmid encoding serotype 6 capsid variant 6.2 and a ITR plasmid containing miniCFTR driven by a minimal CMV promoter into HEK293 cells in a Sartorius Amb250 HT system. At 72h post transfection, cells were lysed by incubating the cells with 10X lysis buffer containing DNase at 37°C for 2h to release intracelluar AAV particles and digest nucleic acids followed by centrifugation to remove cell debris.

### Genome integrity assay by alkaline gel electrophoresis

Affinity chromatography eluates described above were obtained for genomic DNA extraction. Samples were treated with and without DNase I (ThermoFisher AM2224) at 37°C for 30 min to digest the external DNA. Followed by DNAse treatment, QIAamp MinElute Virus Spin Kit (Qiagen 57704) was used to extract encapsidated DNA. In the final step of sample recovery, EB buffer was used to improve the genome recovery. Concentrations of extracted AAV genome were determined by the Nanodrop (Thermo Scientific 840274200) and used accordingly for agarose gel analysis.

Alkaline gel electrophoresis was performed by Owl EasyCast B2 Electrophoresis System (ThermoFisher B2-BP). Agarose gel of 1.2% was prepared from TopVision Agarose Tablets (ThermoFisher R2801) in agarose dissolving buffer (50 mM NaCl, 1 mM EDTA, pH 8.0). The solidified agarose gel was equilibrated the pH in electrophoresis buffer (50 mM NaOH, 1 mM EDTA) for 1 to 1.5 hours. The equilibrated gel was set up in electrophoresis tank and submerged in electrophoresis buffer. One hundred and fifty ng extracted AAV genomic DNA was loaded containing with agarose gel loading dye (Thermo Scientific J62157AB). DNA ladder (ThermoFisher SM0311) was loaded with 500 ng/well with agarose gel loading dye. The electrophoresis was performed at 75V for 3.5 hours at 2-5°C. The gel was neutralized in gel neutralizing buffer (0.5 M Tris-HCl pH 7.5; 1:1 dilution from Corning 46-030-CM) for 30 min and stained with 1:10,000 dilution of SYBR Gold (ThermoFisher S11494) in 1x TBE Buffer (10x dilution from 10x TBE buffer; ThermoFisher 15581044) for 30 min. Gel was imaged with BioRad’s ChemiDoc Gel Imaging System and analyzed with BioRad’s ImageLab Software.

### Mouse AAV Study

Male C57BL/6 mice age 8–10 weeks were injected via the tail vein or via oropharyngeal aspiration with AAVs expressing tdTomato driven by the CAG promoter (n=8 per AAV). The injection administered for each of the serotypes was the same volume as the one used for the serotype with the lowest titer. Control animals were injected with 1× phosphate-buffered saline. Animals were observed for two weeks until study end date at Day 15. Mouse lungs were perfused and dissected in n=4 animals and mouse tracheae were dissected in n=4 animals under sterile conditions. All tissues were immediately fixed in 10% normal buffered formalin for 18–24 h at room temperature then transferred to 70% ethanol and kept at 4°C until paraffin embedding. 5-μm sections were baked and deparaffinized using standard procedures prior to antibody staining.

### Biodistribution study design in NHPs

#### General Study Design

A single dose intravenous injection biodistribution study of AAVs was performed in female naive cynomolgus monkeys with a 21 day observation period at Labcorp.

A single dose of 1 × 10^14^ vg/kg of AAV-173CMV-mCherry (>98% purity) were administered via intravenous (slow bolus) injection on Day 1 to three groups (n = 2) of female naïve cynomolgus monkeys. One group of cynomolgus monkeys (n = 2) was administered 5 mL/kg of TFF3 buffer and served as controls. The monkeys were obtained from Envigo (Alice, Texas).

Clinical observations, body weight measurements and food consumption determinations were performed for all groups. All animals were anesthetized with sodium pentobarbital and exsanguinated on Day 22 of the dosing phase. Macroscopic examinations and organ weight determinations were performed for all groups. Microscopic examinations were conducted on a standard list of organs and tissues from all animals. The following tissues were collected for biodistribution analysis: bone marrow (from femur), brain (frontal cortex), cervix (to be included with the uterus), diaphragm, dorsal root ganglion (one cervical [both sides], one thoracic [both sides], and one lumbar [both sides]), eye (left), gallbladder (apex), gastrointestinal tract (duodenum, ileum, jejunum, and colon), heart (piece of apex), kidney (portion of left kidney pole; cortex only), liver (part of left late**r**al lobe), lung (samples from left cranial, left middle, and left caudal), ovary (left), oviduct (left), pancreas (piece of tail), quadriceps muscle (left), salivary gland (mandibular; left), spinal cord (cervical, thoracic, and lumbar), spleen, trachea, and uterus (included with the cervix). Blocks were created from the following organs for immunohistochemistry and/or in situ hybridization: lung (samples from right cranial, right middle, and right caudal), trachea, liver, duodenum, pancreas, gallbladder, heart, dorsal root ganglion (one cervical [both sides], one thoracic [both sides], and one lumbar [both sides]), and nasal cavity.

#### Quantitative determination of viral DNA in cynomolgus monkey whole blood and tissues by qPCR

DNA purification was performed using the DNeasy 96 Blood and Tissue Kit according to the manufacturer instructions (DNeasy Blood and Tissue Handbook, 2006). Approximately 1 cm^3^ of tissues were used for DNA purification. In the case of Bone Marrow, the pellets present on the bottom of the tube were used. All the samples were shaken 30 seconds at 24 Hz with the TissueLyzer II with 400 μL proteinase K-Buffer ATL solution (10% proteinase K). Lysate was transferred in the microtubes provided with the kit and incubated overnight at 56°C on a thermoshaker. After the incubation, racks were shaken vigorously up and down for 15 seconds and centrifuged. 6 μL of RNase A (100 mg/ml) were added to each sample. Tubes were shaken and incubated 5 minutes at room temperature. Tubes were centrifuged again (centrifuge stopped when reaching 3000 rpm). 600 μL of Buffer AL-ethanol were added and the protocol was continued as described in the DNeasy Blood and Tissue Handbook, 2006 (point 7 page 38). DNA was eluted with 2×100 μL of buffer AE. All DNA samples were quantified by Nanodrop and stored at 4°C.

After DNA purification, all eluted DNAs were quantified by using the ThermoFisher Nanodrop 8000c. The Nanodrop was blanked with eluate buffer from the respective DNA purification kits and purified DNA was quantified at 260 nm for each sample. Concentrations were reported as ng/μL of DNA. The minimal targeted measured concentration on Nanodrop was 12.5; Eluates were normalized with nuclease free water to reach a concentration of 12.5 ng/μL per sample based on the Nanodrop result using Qiagility robot. qPCR was performed in a 384-well plate using the ViiA 7 Fast Real-time qPCR system.

Amplicons spanning the mCherry gene and the RRP30 gene were used to, respectively, measure vector genomes (Vg) and host genomic DNA in a quantitative polymerase chain reaction (qPCR) assay run in duplex. The following primers and probes were used from Integrated DNA Technologies: mCherry probe: 5’-/56-FAM/ACG GCA GCA /ZEN/AGG CCT ACG TG/3IABkFQ/-3’ and mCherry primers 5’-GTA GTC GGG AAT ATC GGC GG-3’ and 5’-GGC AGA CCT TAC GAG GGA AC-3’, mfRPP30 probe 5’/5HEX/GCG GGA GTG /ZEN/TTG GAA CAA TATTAT CCA /3IABkFQ/-3’ mfRPP30 primers 5’-AGG CTC AGA GAA GGA GGA GG-3’ and 5’-ACC GTG TAT GTC CCC ACC TA-3’

#### Immunohistochemistry on tissue sections

Immunohistochemistry staining for mCherry including the deparaffinization and antigen retrieval steps, were performed on a Ventana Discovery XT autostainer using standard Ventana Discovery XT reagents (Ventana, Indianapolis, IN). Slides were deparaffinized then submitted to heat-induced antigen retrieval by covering them with Cell Conditioning 1 (CC1/pH8) solution according to the standard Ventana retrieval protocol. Slides were incubated with the primary antibody (rabbit monoclonal anti-mCherry antibody clone 1C51 at 5.0 ug/mL) or a non-immune isotype-matched control (Rabbit monoclonal IgG2A clone 20102 at 5.0 ug/ml) for one hour. Visualization was obtained by incubation with the appropriate Ventana Discovery OmniMap HRP reagent followed by Ventana Discovery ChromoMap 3,3’-Diaminobenzidine (DAB). Counterstaining was performed using Ventana Hematoxylin and Ventana Bluing reagent for 4 minutes each. Slides were dehydrated, cleared and coverslipped with a synthetic mounting medium.

#### In situ hybridization

In situ hybridization analysis of tissue sections from formalin fixed paraffin embedded blocks was conducted to detect AAV antisense (AS) and sense (S) sequences as well as Hs-PPIB (peptidylprolyl isomerase B [cyclophilin B]; positive control and tissue quality control) and DapB (negative control) genes using reagents and equipment supplied by Advanced Cell Diagnostics (ACDBio, Hayward, CA, USA) and Ventana Medical Systems (Roche, Tuscon AZ, USA). The following ISH RNAscope® probes were designed by ACDBio and do not cross hybridize to the human genome: Hs-PPIB, DapB, and mCherry. The FOXJ1, SCGB1A1, and KRT5 conterstaining probes were intended to identify specific cell types with AAV transduction. Positive PPIB and negative DapB control probe sets were included to ensure mRNA quality and specificity, respectively. The hybridization method followed protocols established by ACDBio and Ventana systems using a 3,3’-Diaminobenzidine (DAB) chromogen. Briefly, 4-5 µm sections were baked at 60°C for 60 minutes and used for hybridization. The deparaffinization and rehydration protocol was performed using a Sakura Tissue-Tek DR5 stainer (Sakura Finetek USA, Torrance, CA) with the following steps: three times xylene for 3 minutes each, two times 100% alcohol for 3 minutes, and air dried for 5 minutes. Off-line manual pretreatment in 1X retrieval buffer at 98°C to 104°C was performed for 60 minutes. Optimization was performed by first evaluating PPIB and DapB hybridization signal and subsequently using the same conditions for all slides. Following pretreatment, the slides were transferred to a Ventana Ultra autostainer to complete the ISH procedure, including protease pretreatment, hybridization at 43°C for 2 hours followed by amplification, and detection with horseradish peroxidase and hematoxylin counter stain.

## Statistical analysis

No statistical methods were used to predetermine sample size. The experiments were not randomized. The investigators were not blinded to allocation during experiments and outcome assessment. The standard error of the mean was calculated from the mean of at least three independent HBEC cultures. The Student’s t-test (unpaired, two-tailed) was used to compare data between groups, with a P value of less than 0.05 considered significant. A one-way ANOVA and post-hoc Tukey’s multiple comparisons test was conducted to determine the rescue of CFTR activity in CF HBECs.

## Author Contributions

L.W.P., C.S., R.W., and A.L. conceived the idea and designed the studies. C.S., B.T. and A.D. performed the structure guided engineering of the CFTR minigenes. J.S. packaged CFTR minigenes in lentiviral vector and L.R., E.S. and R.V.M. tested CFTR minigenes in cell-based assays. A.N. developed and performed LC-MS method for CFTR peptide quantification. A.L., L.A., H.W., M.B., G.T. performed packaging, purification and quality analysis of AAV vectors. R.C.W. performed mouse studies and C.Q. performed IHC on mouse tissues. R.S., D.I.A., and M.M., performed and analyzed IHC on NHP tissues and K.K. and I.W. performed viral genome quantification on NHP tissues. L.W.P. wrote the manuscript with review and support from all authors.

## Acknowledgements

We would like to thank Aron Jaffe, Agostino Cirillo, Hilmar Ebersbach, Deborah Scheffer, Michael Bardroff, Bridget Stuart and Keith Mansfield for project input and guidance. We would also like to thank Jeff Li and Valentina Kemp for help with in vitro assay development, as well as Kristie Wetzel and Paola Capodieci for help with tissue processing and histology. Finally, we greatly appreciate the support from David Rowlands, Timothy Benson, Tewis Bouwmeester and Jay Bradner through the course of this work.

## Supplemental Figures

**Figure S1:**
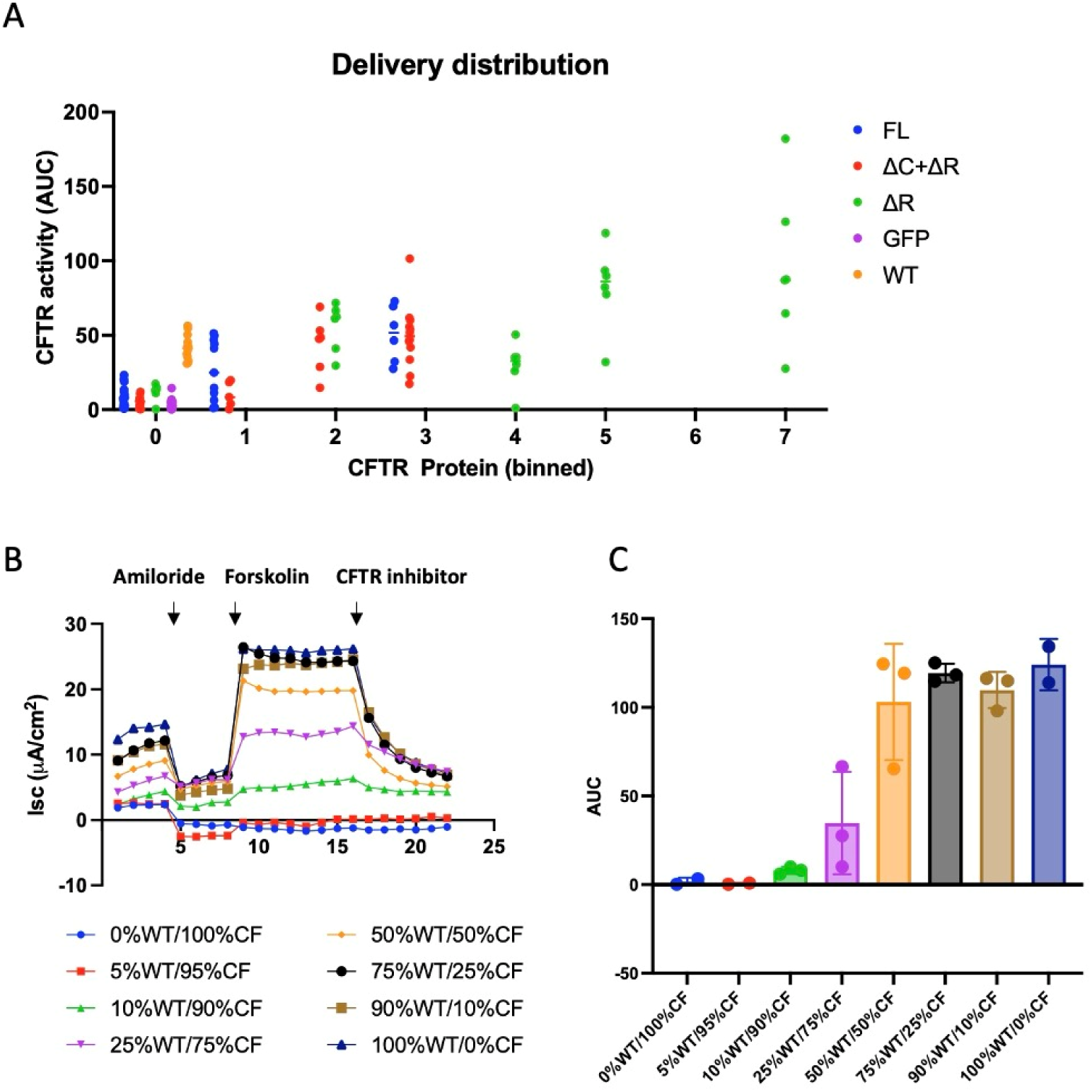
CFTR activity and peptide analysis in Class 1 CF human bronchial epithelial cells (HBECs) following lentiviral delivery of CFTR constructs or reintroduction of non-CF cells to culture. **(A)** One hour after seeding, HBECs were treated with lentivirus expressing full length (FL) CFTR or CFTR minigenes ΔR+ΔC and ΔR. After three weeks of differentiation, CFTR activity was measured by transepithelial chloride conductance (TECC) assay and protein was measured by LC-MS. Given the variable distribution of protein delivered with each lentivirus, samples were binned by protein levels and conditions in which more than one construct was represented and measured LC-MS peptide was greater than that measured in wildtype samples (bins 1-3) were analyzed for **Fig 2D**. (**B,C**) At seeding, non-CF and CF donor HBECs were mixed in the outlined percentages and three weeks later TECC analysis was performed. The average short circuit current (Isc; **B**) and area under the curve (AUC; **C**) in response to CFTR stimulation and inhibition is plotted for each condition (n=2 or 3 wells per condition).

**Figure S2:**
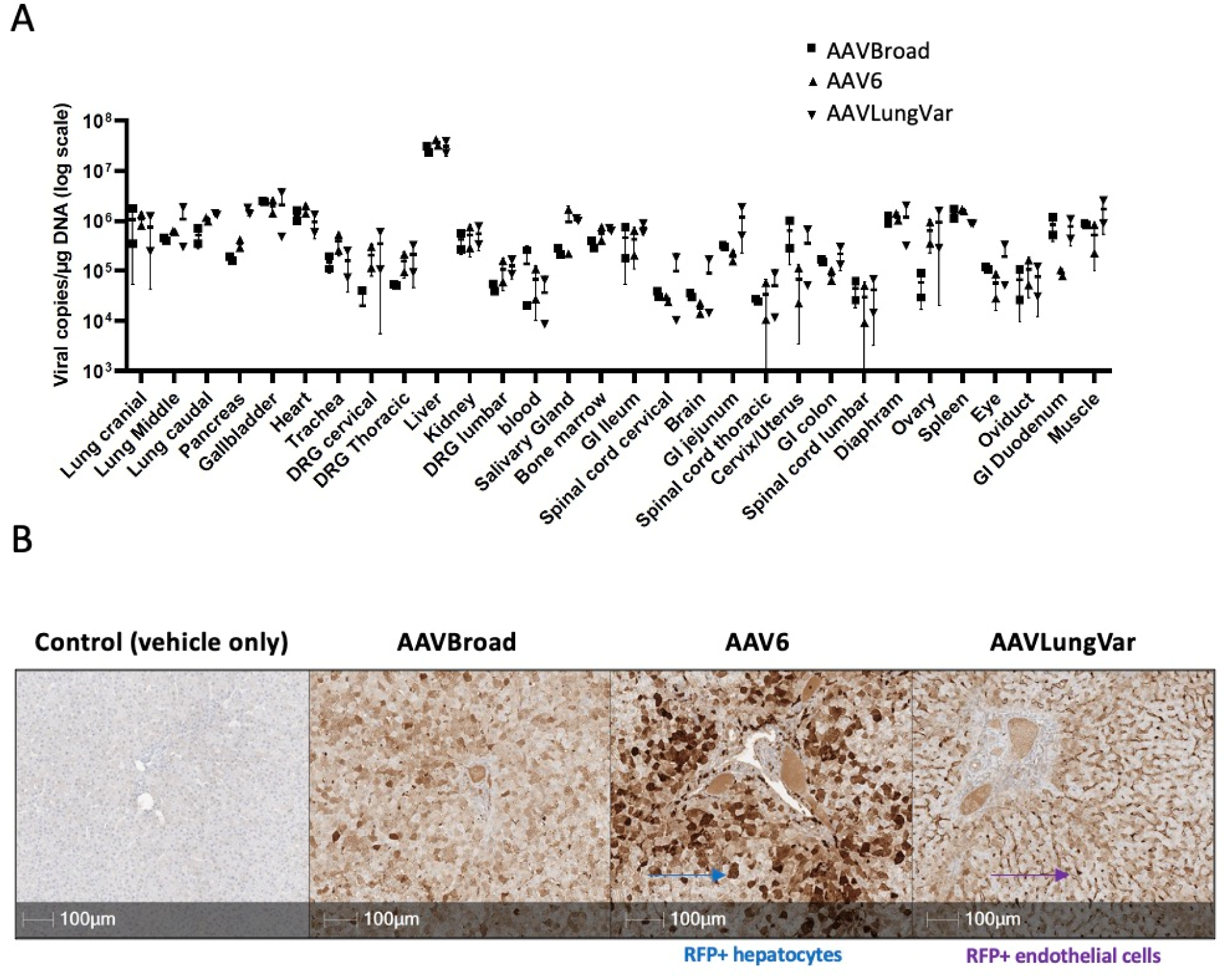
AAV capsid biodistribution across tissues and liver transduction in NHP three weeks after intravenous (IV) delivery at ∼1 × 10^14^ vg/kg. **A** Viral genome DNA was measured in listed tissues (n=2 biological replicates per AAV). **B** Liver transduction and expression of 173CMV-mCherry was assessed by IHC using an RFP antibody (Invitrogen) demonstrating transduction of different cell types by different capsids.

**Figure S3:**
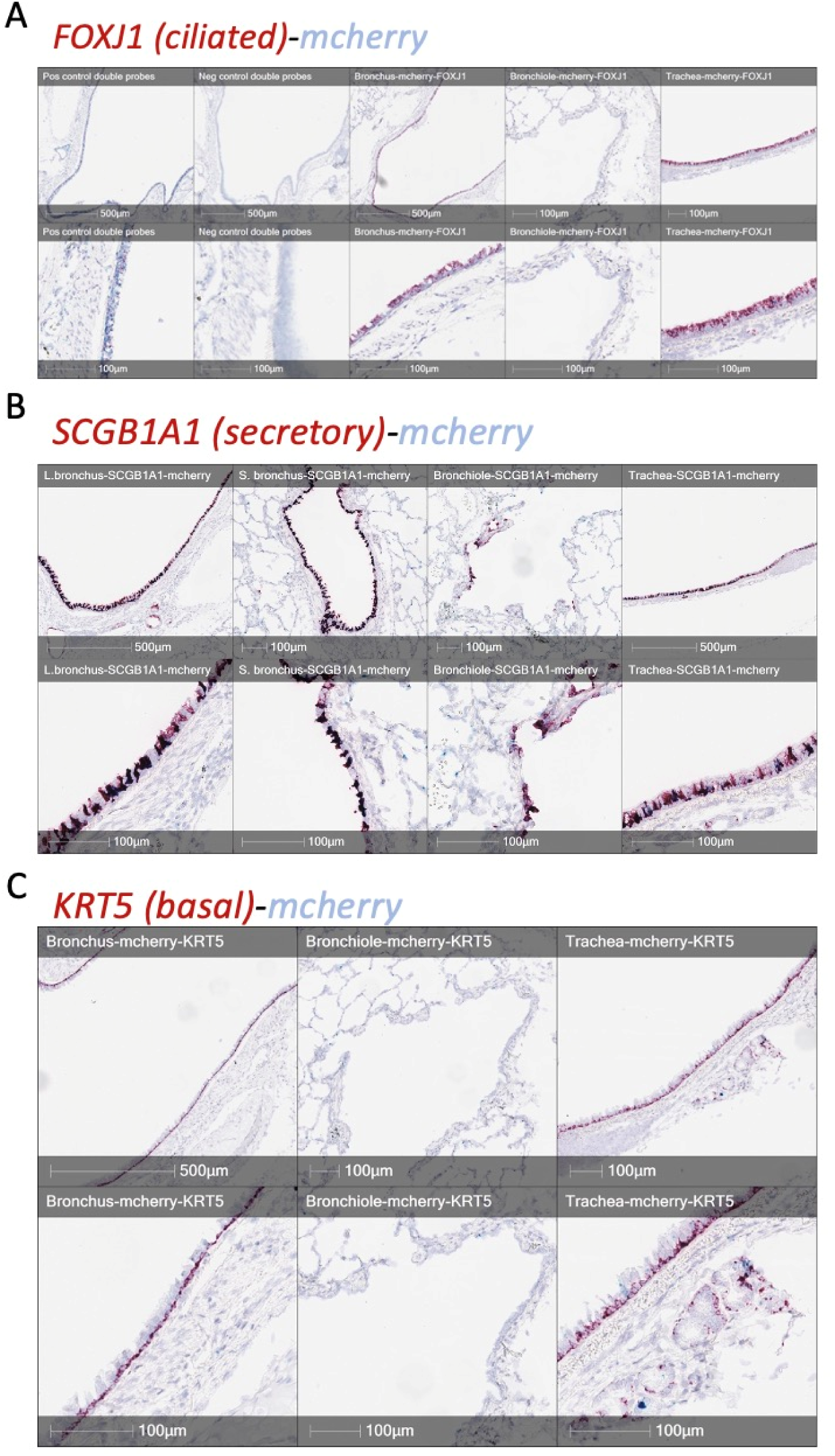
Transduction of proximal airway cell types by AAVLungVar was assessed by RNA *in situ* hybridization (RNAscope) of mcherry reporter and cell type markers. Probes designed to recognize *mcherry* mRNA were combined with probes to label ciliated cells (**A**; FOXJ1), secretory cells (**B**; SCGB1A1) and basal cells (**C**; KRT5).

## References

Alton, E. W., Armstrong, D. K., Ashby, D., Bayfield, K. J., Bilton, D., Bloomfield, E. V, Boyd, A. C., Brand, J., Buchan, R., Calcedo, R., Carvelli, P., Chan, M., Cheng, S. H., Collie, D. S., Cunningham, S., Davidson, H. E., Davies, G., Davies, J. C., Davies, L. A.,… Wolstenholme-Hogg, P. (2016). A randomised, double-blind, placebo-controlled trial of repeated nebulisation of non-viral cystic fibrosis transmembrane conductance regulator (CFTR) gene therapy in patients with cystic fibrosis. Efficacy and Mechanism Evaluation, 3(5), 1–210. 10.3310/EME03050

Al-Zaidy, S. A., Kolb, S. J., Lowes, L., Alfano, L. N., Shell, R., Church, K. R., Nagendran, S., Sproule, D. M., Feltner, D. E., Wells, C., Ogrinc, F., Menier, M., L’Italien, J., Arnold, W. D., Kissel, J. T., Kaspar, B. K., & Mendell, J. R. (2019). AVXS-101 (Onasemnogene Abeparvovec) for SMA1: Comparative Study with a Prospective Natural History Cohort. Journal of Neuromuscular Diseases, 6(3), 307–317. 10.3233/JND-190403

Batista, A. R., King, O. D., Reardon, C. P., Davis, C., Shankaracharya, Philip, V., Gray-Edwards, H., Aronin, N., Lutz, C., Landers, J., & Sena-Esteves, M. (2020). Ly6a Differential Expression in Blood–Brain Barrier Is Responsible for Strain Specific Central Nervous System Transduction Profile of AAV-PHP.B. Https://Home.Liebertpub.Com/Hum, 31(1–2), 90–102. 10.1089/HUM.2019.186

Carneiro, A., Lee, H., Lin, L., Van Haasteren, J., & Schaffer, D. V. (2020). Novel Lung Tropic Adeno-Associated Virus Capsids for Therapeutic Gene Delivery. Human Gene Therapy, 31(17–18), 996–1009. 10.1089/HUM.2020.169

Carroll, T. P., Morales, M. M., Fulmer, S. B., Allen, S. S., Flotte, T. R., Cutting, G. R., & Gugginot, W. B. (1995). Alternate Translation Initiation Codons Can Create Functional Forms of Cystic Fibrosis Transmembrane Conductance Regulator*. In THE JOURNAL OF BIOLOGICAL CHEMISTRY (Vol. 270, Issue 20).

Cebotaru, L., Woodward, O., Cebotaru, V., & Guggino, W. B. (2013). Transcomplementation by a Truncation Mutant of Cystic Fibrosis Transmembrane Conductance Regulator (CFTR) Enhances ΔF508 Processing through a Biomolecular Interaction. The Journal of Biological Chemistry, 288(15), 10505. 10.1074/JBC.M112.420489

Cheng, Q., Wei, T., Farbiak, L., Johnson, L. T., Dilliard, S. A., & Siegwart, D. J. (2020). Selective organ targeting (SORT) nanoparticles for tissue-specific mRNA delivery and CRISPR–Cas gene editing. Nature Nanotechnology 2020 15:4, 15(4), 313–320. 10.1038/s41565-020-0669-6

Choi, J. H., Yu, N. K., Baek, G. C., Bakes, J., Seo, D., Nam, H. J., Baek, S. H., Lim, C. S., Lee, Y. S., & Kaang, B. K. (2014). Optimization of AAV expression cassettes to improve packaging capacity and transgene expression in neurons. Molecular Brain, 7(1). 10.1186/1756-6606-7-17

Cooney, A. L., McCray, P. B., & Sinn, P. L. (2018). Cystic Fibrosis Gene Therapy: Looking Back, Looking Forward. Genes, 9(11). 10.3390/GENES9110538

Das, A., Vijayan, M., Walton, E. M., Stafford, V. G., Fiflis, D. N., & Asokan, A. (2022a). Epigenetic Silencing of Recombinant Adeno-associated Virus Genomes by NP220 and the HUSH Complex. Journal of Virology, 96(4). 10.1128/jvi.02039-21

Das, A., Vijayan, M., Walton, E. M., Stafford, V. G., Fiflis, D. N., & Asokan, A. (2022b). Epigenetic Silencing of Recombinant Adeno-associated Virus Genomes by NP220 and the HUSH Complex. Journal of Virology, 96(4). 10.1128/jvi.02039-21

Dong, J. Y., Fan, P. D., & Frizzell, R. A. (1996). Quantitative analysis of the packaging capacity of recombinant adeno-associated virus. Human Gene Therapy, 7(17), 2101–2112. 10.1089/HUM.1996.7.17-2101

Duncan, G. A., Kim, N., Colon-Cortes, Y., Rodriguez, J., Mazur, M., Birket, S. E., Rowe, S. M., West, N. E., Livraghi-Butrico, A., Boucher, R. C., Hanes, J., Aslanidi, G., & Suk, J. S. (2018). An Adeno-Associated Viral Vector Capable of Penetrating the Mucus Barrier to Inhaled Gene Therapy. Molecular Therapy Methods and Clinical Development, 9, 296–304. 10.1016/J.OMTM.2018.03.006/ATTACHMENT/60C2343C-0F9C-418D-A6B9-5CEDFF2FD333/MMC2.PDF

Excoffon, K. J. D. A., Koerber, J. T., Dickey, D. D., Murtha, M., Keshavjee, S., Kaspar, B. K., Zabner, J., & Schaffer, D. V. (2009). Directed evolution of adeno-associated virus to an infectious respiratory virus. Proceedings of the National Academy of Sciences of the United States of America, 106(10), 3865–3870. 10.1073/PNAS.0813365106/SUPPL_FILE/0813365106SI.PDF

Excoffon, K. J. D. A., Smith, M. D., Falese, L., Schulingkamp, R., Lin, S., Mahankali, M., Narayan, P. K. L., Glatfelter, M. R., Limberis, M. P., Yuen, E., & Kolbeck, R. (2024). Inhalation of SP-101 Followed by Inhaled Doxorubicin Results in Robust and Durable hCFTRΔR Transgene Expression in the Airways of Wild-Type and Cystic Fibrosis Ferrets. Human Gene Therapy. 10.1089/HUM.2024.064

Fischer, A. C., Smith, C. I., Cebotaru, L., Zhang, X., Askin, F. B., Wright, J., Guggino, S. E., Adams, R. J., Flotte, T., & Guggino, W. B. (2007). Expression of a truncated cystic fibrosis transmembrane conductance regulator with an AAV5-pseudotyped vector in primates. Molecular Therapy : The Journal of the American Society of Gene Therapy, 15(4), 756–763. 10.1038/SJ.MT.6300059

Furusho, T., Das, R., Hakui, H., Sairavi, A., Adachi, K., Galbraith-Liss, M. S., Rajagopal, P., Horikawa, M., Luo, S., Li, L., Yamada, K., Andeen, N., Dissen, G. A., & Nakai, H. (2024). Enhancing gene transfer to renal tubules and podocytes by context-dependent selection of AAV capsids. Nature Communications, 15(1), 10728. 10.1038/s41467-024-54475-9

Goertsen, D., Goeden, N., Flytzanis, N. C., & Gradinaru, V. (2022). Targeting the lung epithelium after intravenous delivery by directed evolution of underexplored sites on the AAV capsid. Molecular Therapy Methods and Clinical Development, 26, 331–342. 10.1016/J.OMTM.2022.07.010/ATTACHMENT/462DFD21-7994-4228-ACF1-6474FB768E50/MMC2.PDF

Gonzalez-Sandoval, A., Pekrun, K., Tsuji, S., Zhang, F., Hung, K. L., Chang, H. Y., & Kay, M. A. (2023). The AAV capsid can influence the epigenetic marking of rAAV delivered episomal genomes in a species dependent manner. Nature Communications, 14(1). 10.1038/s41467-023-38106-3

Halbert, C. L., Allen, J. M., & Miller, A. D. (2001). Adeno-Associated Virus Type 6 (AAV6) Vectors Mediate Efficient Transduction of Airway Epithelial Cells in Mouse Lungs Compared to That of AAV2 Vectors. Journal of Virology, 75(14), 6615–6624. 10.1128/jvi.75.14.6615-6624.2001

Harvey, B. G., Leopold, P. L., Hackett, N. R., Grasso, T. M., Williams, P. M., Tucker, A. L., Kaner, R. J., Ferris, B., Gonda, I., Sweeney, T. D., Ramalingam, R., Kovesdi, I., Shak, S., & Crystal, R. G. (1999). Airway epithelial CFTR mRNA expression in cystic fibrosis patients after repetitive administration of a recombinant adenovirus. The Journal of Clinical Investigation, 104(9), 1245–1255. 10.1172/JCI7935

Heijerman, H. G. M., McKone, E. F., Downey, D. G., Van Braeckel, E., Rowe, S. M., Tullis, E., Mall, M. A., Welter, J. J., Ramsey, B. W., McKee, C. M., Marigowda, G., Moskowitz, S. M., Waltz, D., Sosnay, P. R., Simard, C., Ahluwalia, N., Xuan, F., Zhang, Y., Taylor-Cousar, J. L.,… Majoor, C. (2019). Efficacy and safety of the elexacaftor/tezacaftor/ivacaftor combination regimen in people with cystic fibrosis homozygous for the F508del mutation: a double-blind, randomised, phase 3 trial. *Lancet (London*, England), 394(10212), 1940. 10.1016/S0140-6736(19)32597-8

Hirsch, M. L., Wolf, S. J., & Samulski, R. J. (2016). Delivering Transgenic DNA Exceeding the Carrying Capacity of AAV Vectors. *Methods in Molecular Biology (Clifton*, N.J.), 1382(1), 21. 10.1007/978-1-4939-3271-9_2

Hordeaux, J., Lamontagne, R. J., Song, C., Buchlis, G., Dyer, C., Buza, E. L., Ramezani, A., Wielechowski, E., Greig, J. A., Chichester, J. A., Bell, P., & Wilson, J. M. (2024). High-dose systemic adeno-associated virus vector administration causes liver and sinusoidal endothelial cell injury. Molecular Therapy, 32(4), 952–968. 10.1016/j.ymthe.2024.02.002

Hubert, D., Chiron, R., Camara, B., Grenet, D., Prévotat, A., Bassinet, L., Dominique, S., Rault, G., Macey, J., Honoré, I., Kanaan, R., Leroy, S., Desmazes Dufeu, N., & Burgel, P. R. (2017). Real-life initiation of lumacaftor/ivacaftor combination in adults with cystic fibrosis homozygous for the Phe508del CFTR mutation and severe lung disease. Journal of Cystic Fibrosis : Official Journal of the European Cystic Fibrosis Society, 16(3), 388–391. 10.1016/J.JCF.2017.03.003

Johnson, L. G., Olsen, J. C., Sarkadi, B., Moore, K. L., Swanstrom, R., & Boucher, R. C. (1992). Efficiency of gene transfer for restoration of normal airway epithelial function in cystic fibrosis. Nature Genetics, 2(1), 21–25. 10.1038/NG0992-21

Keating, D., Marigowda, G., Burr, L., Daines, C., Mall, M. A., McKone, E. F., Ramsey, B. W., Rowe, S. M., Sass, L. A., Tullis, E., McKee, C. M., Moskowitz, S. M., Robertson, S., Savage, J., Simard, C., Van Goor, F., Waltz, D., Xuan, F., Young, T., & Taylor-Cousar, J. L. (2018). VX-445-Tezacaftor-Ivacaftor in Patients with Cystic Fibrosis and One or Two Phe508del Alleles. The New England Journal of Medicine, 379(17), 1612–1620. 10.1056/NEJMOA1807120

Konkimalla, A., Elmore, Z., Konishi, S., Macadlo, L., Katsura, H., Tata, A., Asokan, A., & Tata, P. R. (2023a). Efficient Adeno-associated Virus-mediated Transgenesis in Alveolar Stem Cells and Associated Niches. American Journal of Respiratory Cell and Molecular Biology, 69(3), 255–265. 10.1165/rcmb.2022-0424MA

Konkimalla, A., Elmore, Z., Konishi, S., Macadlo, L., Katsura, H., Tata, A., Asokan, A., & Tata, P. R. (2023b). Efficient Adeno-associated Virus-mediated Transgenesis in Alveolar Stem Cells and Associated Niches. American Journal of Respiratory Cell and Molecular Biology, 69(3), 255–265. 10.1165/RCMB.2022-0424MA

Limberis, M. P., Vandenberghe, L. H., Zhang, L., Pickles, R. J., & Wilson, J. M. (2009). Transduction efficiencies of novel AAV vectors in mouse airway epithelium in vivo and human ciliated airway epithelium in vitro. Molecular Therapy, 17(2), 294–301. 10.1038/mt.2008.261

Liu, F., Zhang, Z., Csanády, L., Gadsby, D. C., & Chen, J. (2017). Molecular Structure of the Human CFTR Ion Channel. Cell, 169(1), 85–95.e8. 10.1016/J.CELL.2017.02.024

McBennett, K. A., Davis, P. B., & Konstan, M. W. (2022). Increasing life expectancy in cystic fibrosis: Advances and challenges. Pediatric Pulmonology, 57(Suppl 1), S5. 10.1002/PPUL.25733

McCarron, A., Cmielewski, P., Drysdale, V., Parsons, D., & Donnelley, M. (2023). Effective viral-mediated lung gene therapy: is airway surface preparation necessary? Gene Therapy, 30(6), 469–477. 10.1038/S41434-022-00332-7

Middleton, P. G., Mall, M. A., Dřevínek, P., Lands, L. C., McKone, E. F., Polineni, D., Ramsey, B. W., Taylor-Cousar, J. L., Tullis, E., Vermeulen, F., Marigowda, G., McKee, C. M., Moskowitz, S. M., Nair, N., Savage, J., Simard, C., Tian, S., Waltz, D., Xuan, F.,… Jain, R. (2019). Elexacaftor–Tezacaftor–Ivacaftor for Cystic Fibrosis with a Single Phe508del Allele. New England Journal of Medicine, 381(19), 1809–1819. 10.1056/NEJMOA1908639/SUPPL_FILE/NEJMOA1908639_DATA-SHARING.PDF

Montoro, D. T., Haber, A. L., Biton, M., Vinarsky, V., Lin, B., Birket, S. E., Yuan, F., Chen, S., Leung, H. M., Villoria, J., Rogel, N., Burgin, G., Tsankov, A. M., Waghray, A., Slyper, M., Waldman, J., Nguyen, L., Dionne, D., Rozenblatt-Rosen, O.,… Rajagopal, J. (2018). A revised airway epithelial hierarchy includes CFTR-expressing ionocytes. Nature, 560(7718), 319–324. 10.1038/s41586-018-0393-7

Ostedgaard, L. S., Meyerholz, D. K., Vermeer, D. W., Karpa, P. H., Schneider, L., Sigmund, C. D., & Welsha, M. J. (2011). Cystic fibrosis transmembrane conductance regulator with a shortened R domain rescues the intestinal phenotype of CFTR-/- mice. Proceedings of the National Academy of Sciences of the United States of America, 108(7), 2921–2926. 10.1073/PNAS.1019752108

Ostedgaard, L. S., Rokhlina, T., Karp, P. H., Lashmit, P., Afione, S., Schmidt, M., Zabner, J., Stinski, M. F., Chiorini, J. A., & Welsh, M. J. (2005a). A shortened adeno-associated virus expression cassette for CFTR gene transfer to cystic fibrosis airway epithelia. Proceedings of the National Academy of Sciences of the United States of America, 102(8), 2952–2957. 10.1073/PNAS.0409845102

Ostedgaard, L. S., Rokhlina, T., Karp, P. H., Lashmit, P., Afione, S., Schmidt, M., Zabner, J., Stinski, M. F., Chiorini, J. A., & Welsh, M. J. (2005b). A shortened adeno-associated virus expression cassette for CFTR gene transfer to cystic fibrosis airway epithelia. Proceedings of the National Academy of Sciences of the United States of America, 102(8), 2952–2957. 10.1073/PNAS.0409845102

Ostedgaard, L. S., Zabner, J., Vermeer, D. W., Rokhlina, T., Karp, P. H., Stecenko, A. A., Randak, C., & Welsh, M. J. (2002). CFTR with a partially deleted R domain corrects the cystic fibrosis chloride transport defect in human airway epithelia in vitro and in mouse nasal mucosa in vivo. Proceedings of the National Academy of Sciences of the United States of America, 99(5), 3093–3098. 10.1073/PNAS.261714599

Plasschaert, L. W., MacDonald, K. D., & Moffit, J. S. (2024). Current landscape of cystic fibrosis gene therapy. In Frontiers in Pharmacology (Vol. 15). Frontiers Media SA. 10.3389/fphar.2024.1476331

Plasschaert, L. W., Žilionis, R., Choo-Wing, R., Savova, V., Knehr, J., Roma, G., Klein, A. M., & Jaffe, A. B. (2018). A single-cell atlas of the airway epithelium reveals the CFTR-rich pulmonary ionocyte. Nature, 560(7718), 377–381. 10.1038/s41586-018-0394-6

Potter, R. A., Griffin, D. A., Heller, K. N., Peterson, E. L., Clark, E. K., Mendell, J. R., & Rodino-Klapac, L. R. (2021). Dose-Escalation Study of Systemically Delivered rAAVrh74.MHCK7.micro-dystrophin in the mdx Mouse Model of Duchenne Muscular Dystrophy. Human Gene Therapy, 32(7–8), 375–389. 10.1089/HUM.2019.255

Ramalho, A. S., Beck, S., Meyer, M., Penque, D., Cutting, G. R., & Amaral, M. D. (2002). Five percent of normal cystic fibrosis transmembrane conductance regulator mRNA ameliorates the severity of pulmonary disease in cystic fibrosis. American Journal of Respiratory Cell and Molecular Biology, 27(5), 619–627. 10.1165/RCMB.2001-0004OC

Riordan, J. R., Rommens, J. M., Kerem, B. S., Alon, N. O. A., Rozmahel, R., Grzelczak, Z., Zielenski, J., Lok, S. I., Plavsic, N., Chou, J. L., Drumm, M. L., Iannuzzi, M. C., Collins, F. S., & Tsui, L. C. (1989). Identification of the cystic fibrosis gene: cloning and characterization of complementary DNA. Science (New York, N.Y.), 245(4922), 1066–1073. 10.1126/SCIENCE.2475911

Rowe, S. M., Zuckerman, J. B., Dorgan, D., Lascano, J., McCoy, K., Jain, M., Schechter, M. S., Lommatzsch, S., Indihar, V., Lechtzin, N., McBennett, K., Callison, C., Brown, C., Liou, T. G., MacDonald, K. D., Nasr, S. Z., Bodie, S., Vaughn, M., Meltzer, E. B., & Barbier, A. J. (2023). Inhaled mRNA therapy for treatment of cystic fibrosis: Interim results of a randomized, double-blind, placebo-controlled phase 1/2 clinical study. Journal of Cystic Fibrosis : Official Journal of the European Cystic Fibrosis Society, 22(4), 656–664. 10.1016/J.JCF.2023.04.008

Sato, Y., Mustafina, K. R., Luo, Y., Martini, C., Thomas, D. Y., Wiseman, P. W., & Hanrahan, J. W. (2021). Nonspecific binding of common anti-CFTR antibodies in ciliated cells of human airway epithelium. Scientific Reports, 11(1). 10.1038/S41598-021-02420-X

Steines, B., Dickey, D. D., Bergen, J., Excoffon, K. J. D. A., Weinstein, J. R., Li, X., Yan, Z., Abou Alaiwa, M. H., Shah, V. S., Bouzek, D. C., Powers, L. S., Gansemer, N. D., Ostedgaard, L. S., Engelhardt, J. F., Stoltz, D. A., Welsh, M. J., Sinn, P. L., Schaffer, D. V., & Zabner, J. (2016). CFTR gene transfer with AAV improves early cystic fibrosis pig phenotypes. JCI Insight, 1(14). 10.1172/JCI.INSIGHT.88728

Steines, B., Dickey, D. D., Bergen, J., JDA Excoffon, K., Weinstein, J. R., Li, X., Yan, Z., Abou Alaiwa, M. H., Shah, V. S., Bouzek, D. C., Powers, L. S., Gansemer, N. D., Ostedgaard, L. S., Engelhardt, J. F., Stoltz, D. A., Welsh, M. J., Sinn, P. L., Schaffer, D. V, Zabner, J., & Carver, L. A. (2016). CFTR gene transfer with AAV improves early cystic fibrosis pig phenotypes. Anatomy and Cell Biology, 1(14). 10.1172/jci.insight.88728

Van Goor, F., Hadida, S., Grootenhuis, P. D. J., Burton, B., Stack, J. H., Straley, K. S., Decker, C. J., Miller, M., McCartney, J., Olson, E. R., Wine, J. J., Frizzell, R. A., Ashlock, M., & Negulescu, P. A. (2011). Correction of the F508del-CFTR protein processing defect in vitro by the investigational drug VX-809. Proceedings of the National Academy of Sciences of the United States of America, 108(46), 18843–18848. 10.1073/pnas.1105787108

Wagner, J. A., Nepomuceno, I. B., Messner, A. H., Moran, M. L., Batson, E. P., Dimiceli, S., Brown, B. W., Desch, J. K., Norbash, A. M., Conrad, C. K., Guggino, W. B., Flotte, T. R., Wine, J. J., Carter, B. J., Reynolds, T. C., Moss, R. B., & Gardner, P. (2002). A phase II, double-blind, randomized, placebo-controlled clinical trial of tgAAVCF using maxillary sinus delivery in patients with cystic fibrosis with antrostomies. Human Gene Therapy, 13(11), 1349–1359. 10.1089/104303402760128577

Wainwright, C. E., Elborn, J. S., Ramsey, B. W., Marigowda, G., Huang, X., Cipolli, M., Colombo, C., Davies, J. C., De Boeck, K., Flume, P. A., Konstan, M. W., McColley, S. A., McCoy, K., McKone, E. F., Munck, A., Ratjen, F., Rowe, S. M., Waltz, D., & Boyle, M. P. (2015). Lumacaftor–Ivacaftor in Patients with Cystic Fibrosis Homozygous for Phe508del CFTR. New England Journal of Medicine, 373(3), 220–231. 10.1056/nejmoa1409547

Wang, B., Li, J., & Xiao, X. (2000). Adeno-associated virus vector carrying human minidystrophin genes effectively ameliorates muscular dystrophy in mdx mouse model. Proceedings of the National Academy of Sciences of the United States of America, 97(25), 13714–13719. 10.1073/PNAS.240335297

Watakabe, A., Ohtsuka, M., Kinoshita, M., Takaji, M., Isa, K., Mizukami, H., Ozawa, K., Isa, T., & Yamamori, T. (2015). Comparative analyses of adeno-associated viral vector serotypes 1, 2, 5, 8 and 9 in marmoset, mouse and macaque cerebral cortex. Neuroscience Research, 93, 144–157. 10.1016/j.neures.2014.09.002

Wu, H., Zhao, A., Bu, Y., Yang, W., He, L., Zhong, Y., Yao, D., Li, H., & Yin, W. (2024). Tropism of adeno-associated virus serotypes in mouse lungs via intratracheal instillation. Virology Journal, 21(1), 302. 10.1186/s12985-024-02575-9

Wu, Z., Asokan, A., Grieger, J. C., Govindasamy, L., Agbandje-McKenna, M., & Samulski, R. J. (2006). Single Amino Acid Changes Can Influence Titer, Heparin Binding, and Tissue Tropism in Different Adeno-Associated Virus Serotypes. Journal of Virology, 80(22), 11393–11397. 10.1128/jvi.01288-06

Yan, Z., Zak, R., Zhang, Y., Ding, W., Godwin, S., Munson, K., Peluso, R., & Engelhardt, J. F. (2004). Distinct classes of proteasome-modulating agents cooperatively augment recombinant adeno-associated virus type 2 and type 5-mediated transduction from the apical surfaces of human airway epithelia. Journal of Virology, 78(6), 2863–2874. 10.1128/JVI.78.6.2863-2874.2004

Zhang, L., Button, B., Gabriel, S. E., Burkett, S., Yan, Y., Skiadopoulos, M. H., Dang, Y. L., Vogel, L. N., McKay, T., Mengos, A., Boucher, R. C., Collins, P. L., & Pickles, R. J. (2009). CFTR delivery to 25% of surface epithelial cells restores normal rates of mucus transport to human cystic fibrosis airway epithelium. PLoS Biology, 7(7). 10.1371/JOURNAL.PBIO.1000155

Zhang, L. N., Karp, P., Gerard, C. J., Pastor, E., Laux, D., Munson, K., Yan, Z., Liu, X., Godwin, S., Thomas, C. P., Zabner, J., Shi, H., Caldwell, C. W., Peluso, R., Carter, B., & Engelhardt, J. F. (2004). Dual therapeutic utility of proteasome modulating agents for pharmaco-gene therapy of the cystic fibrosis airway. Molecular Therapy : The Journal of the American Society of Gene Therapy, 10(6), 990–1002. 10.1016/J.YMTHE.2004.08.009

Zhang, L., Wang, D., Fischer, H., Fan, P. D., Widdicombe, J. H., Kan, Y. W., & Dong, J. Y. (1998). Efficient expression of CFTR function with adeno-associated virus vectors that carry shortened CFTR genes. Proceedings of the National Academy of Sciences of the United States of America, 95(17), 10158–10163. 10.1073/PNAS.95.17.10158

Zhang, Z., Liu, F., & Chen, J. (2018). Molecular structure of the ATP-bound, phosphorylated human CFTR. Proceedings of the National Academy of Sciences of the United States of America, 115(50), 12757–12762. 10.1073/pnas.1815287115

Zincarelli, C., Soltys, S., Rengo, G., & Rabinowitz, J. E. (2008). Analysis of AAV serotypes 1-9 mediated gene expression and tropism in mice after systemic injection. Molecular Therapy, 16(6), 1073–1080. 10.1038/mt.2008.76

